# Plants cope with fluctuating light by frequency-dependent non-photochemical quenching and cyclic electron transport

**DOI:** 10.1101/2022.02.09.479783

**Authors:** Yuxi Niu, Dušan Lazár, Alfred R. Holzwarth, David M. Kramer, Shizue Matsubara, Fabio Fiorani, Hendrik Poorter, Silvia D. Schrey, Ladislav Nedbal

**Affiliations:** Institute of Bio- and Geosciences/Plant Sciences, Forschungszentrum Jülich, Wilhelm-Johnen-Straße, D-52428 Jülich, Germany; Department of Biophysics, Centre of the Region Haná for Biotechnological and Agricultural Research, Faculty of Science, Palacký University, Šlechtitelů 27, 783 71 Olomouc, Czech Republic; Department of Physics and Astronomy, Faculty of Science, Vrije Universiteit Amsterdam, De Boelelaan 1105, NL-1081 HV Amsterdam, The Netherlands; MSU-DOE Plant Research Laboratory, Michigan State University, East Lansing, MI 48824, USA; Department of Natural Sciences, Macquarie University, North Ryde, Australia; PASTEUR, Department of Chemistry, École Normale Supérieure, Université PSL, Sorbonne Université, CNRS, 24, rue Lhomond, 75005, Paris, France

**Keywords:** cyclic electron transport, frequency analysis, non-photochemical quenching, photosynthetic oscillation, regulation

## Abstract

In natural environments, plants are exposed to rapidly changing light. Maintaining photosynthetic efficiency while avoiding photodamage requires equally rapid regulation of photoprotective mechanisms. We asked what the operation frequency range of regulation is in which plants can efficiently respond to varying light.

Chlorophyll fluorescence, P700, plastocyanin, and ferredoxin responses of wild-type *Arabidopsis thaliana* were measured in oscillating light of various frequencies. We also investigated the *npq1* mutant lacking violaxanthin de-epoxidase, the *npq4* mutant lacking PsbS-protein, and the mutants *crr2-2*, and *pgrl1ab* impaired in different pathways of the cyclic electron transport.

The fastest was the PsbS-regulation responding to oscillation periods longer than 10s. Processes involving violaxanthin de-epoxidase dampened changes of chlorophyll fluorescence in oscillation periods of 2min or longer. Knocking out the PGRL1-PGR5 pathway strongly reduced variations of all monitored parameters, probably due to congestion in the electron transport. Incapacitating the NDH-like pathway only slightly changed the photosynthetic dynamics.

Our observations are consistent with the assumption that non-photochemical quenching in slow light oscillations involves violaxanthin de-epoxidase to produce, presumably, a stationary, non-oscillating level of zeaxanthin. We interpret the observed dynamics of Photosystem I components as being formed in slow light oscillations partially by thylakoid remodeling that modulates the redox rates.

## Introduction

Plants grow in dynamic environments and can thrive also in rapidly fluctuating light (Muller *et al.*, 2001; Külheim *et al.*, 2002; Long *et al.*, 2022). Repeated filling and emptying of the biochemical pools in fluctuating light may cause damage by generating transient imbalances between tightly coupled redox reactions, potentially producing noxious reactive oxygen species (Pospíšil, 2009; Kono *et al.*, 2014; Kono & Terashima, 2014). Plants limit the damage and optimize their photosynthetic performance in fluctuating light by regulatory mechanisms that operate with different activation and deactivation rates which constrain the range of frequencies to which the regulation can effectively respond. Illustrative analogies are the range of audio frequencies that a human ear can hear or visible light range that the human eye can see. Here, we report on the limits of the operation range of photosynthetic non-photochemical quenching regulation and cyclic electron transport in the frequency domain.

Light fluctuations in nature may occur with a wide range of characteristic frequencies. Transient gaps in the upper canopy (Chazdon & Pearcy, 1991), wind-induced canopy movement (Peressotti *et al.*, 2001), and intermittent cloudiness (Knapp & Smith, 1987) give rise to irregular light patterns within canopies (Way & Pearcy, 2012; Smith & Berry, 2013; Kaiser *et al.*, 2018). According to the categorization of light fluctuations in canopy of higher plants established by Smith & Berry (2013), sunfleck was classified as light fluctuations with the periods below 8 min, which is also the upper limit of periods investigated here. The short period limit of 1 s of our experiments was dictated by the capacity of the used instrument to generate fast light oscillations.

The method for investigating irregular or periodic stimulation has long been established in physics (Pintelon & Schoukens, 2012) and the used terminology is summarized here in Tab.1. Based on universal mathematical principles, random fluctuation of light can be reproduced as a superposition of harmonic functions, each determined by the period and amplitude of sinus and cosinus oscillations (Schwartz, 2008). If some of the characteristic elemental oscillations are, for example, too fast and thus outside of the operation range, then, the photosynthetic regulation may be less effective. Similarly, when the operation range is reduced, e.g., by a mutation, one can expect different dynamic responses in elemental oscillating light in the wild-type and in the mutant. The essentials of the frequency-domain analysis are summarized here in *Supporting Information – Introduction* and in further detail available in (Nedbal & Březina, 2002; Nedbal *et al.*, 2005; Nedbal & Lazár, 2021; Lazár *et al.*, 2022). Nedbal & Lazár (2021) looked for the abrupt, qualitative changes in photosynthetic responses to oscillating light that would signal potential limits of operation frequency range in the green alga *Chlorella sorokiniana.* The experiments included light oscillations with periods ranging from 1 ms to 512 s and identified four frequency domains (α_1_, β_1_, α_2_, and β_2_), in which oxygenic photosynthesis exhibited contrasting dynamic features. In the α domains, the amplitudes of the ChlF oscillations were found to decrease in algae when the light modulation was slower. The trend was opposite in the β domains. Two of these domains, α and β, in the range between 1 s and 8 min, that are relevant also in canopies of higher plants are in focus of our investigation here *(Supporting Information – Introduction Fig.SI-1*).

**Tab.1:** Glossary of terms.

| Tab.1: Glossary of terms |  |
| --- | --- |
| *amplitude* | the maximum deviation from a mean; in the case of harmonically modulated light, it is one half of the difference between the light intensity maximum and minimum |
| *angular frequency $\omega$* | the parameter of light modulation or of plant response that defines how many times a full cycle (360° in angle degrees or $2\pi$ in radian units) is completed in a unit of time. It is related to the period T as: $\omega = \frac{2\pi}{T}$ |
| *fluctuating light* | randomly changing light: its intensity pattern thus cannot be predicted |
| *forced oscillation* | a repeating variation pattern of plant photosynthesis that is sustained in time by a *periodic* energy supply, here *harmonically modulated light* |
| *frequency-domain measurement* | the measurement of plant activity applying *harmonically modulated light* with *period/frequency* of the light modulation changing over a wide range; plant response(s) are measured as a function of the light modulation *frequency* |
| *harmonically modulated light* | the light intensity that consists of a variable part that is changing as a harmonic function<br>$a * \sin\left(\frac{2\pi}{T} \cdot t\right) + b * \cos\left(\frac{2\pi}{T} \cdot t\right)$ , where $t$ stands for time, $T$ is the *period*, $\frac{2\pi}{T}$ the *angular frequency* of the light variation, and $a, b$ are parameters that define together the *amplitude* and *phase* of the light modulation. |
| *oscillation* | a repeating variation pattern of light or plant activity |
| *period T* | the time after which a process is repeated, here it applies both to the light modulation as well as to plant response elicited by the modulated excitation light |
| *periodic* | Repeating |
| *oscillation phase* | the parameter defining the progress of a periodic process: e.g., phase 0 is the origin, the phase $\frac{1}{4} \cdot 2\pi$ represents one-quarter of the full cycle, phase $2\pi$ represents the full cycle |
| *resonance* | a phenomenon of the increased *amplitude* of a measured response that occurs when the *frequency* of external forcing agrees with an internal characteristic *frequency* of the examined plant (sometimes called eigen-frequency) |
| *spontaneous oscillation* | the *oscillation* of plant activities that sometimes appear in response to an abrupt change of external conditions; spontaneous oscillations of plants are fading over time |
| *time-domain measurement* | the meaning common in fluorometry or spectroscopy of plants: the plant response to an aperiodic light change; mostly a stepwise dark-to-light or light-to-dark transition, often also the response to a short light flash measured as a function of time |

*Arabidopsis thaliana* is exposed to fluctuating light due to intermittent cloudiness, and by canopy movements in natural mixed plant communities (Mitchell-Olds, 2001; Pantazopoulou *et al.*, 2021). It is a model plant that represents well regulation in higher plants and is available with a wide range of well-defined mutations. We used elemental oscillating light with various periods to study its response to light fluctuations in the laboratory. Here, we focus on the responses of various components in the photosynthetic electron transport chain to fluctuating light (Fig.1A), in particular, on non-photochemical quenching (NPQ) and cyclic electron transport (CET) and their dynamics in fluctuating light (Holzwarth *et al.*, 2009; Ruban, 2016; Yamamoto & Shikanai, 2019; Murchie & Ruban, 2020). The rapidly reversible component of NPQ protects the plant from transient excess light by sensing the high-light-induced acidification of the thylakoid lumen and reducing the flow of excitation energy to PSII reaction centers (Demmig-Adams *et al.*, 2014; Ware *et al.*, 2015). This NPQ involves the protonation of PsbS protein (Li *et al.*, 2002; Roach & Krieger-Liszkay, 2012; Ruban, 2016) and the de-epoxidation of violaxanthin to zeaxanthin by the violaxanthin de-epoxidase enzyme (VDE) (Fig.1B) (Gilmore, 1997; Jahns & Holzwarth, 2012).

**Fig.1A.**
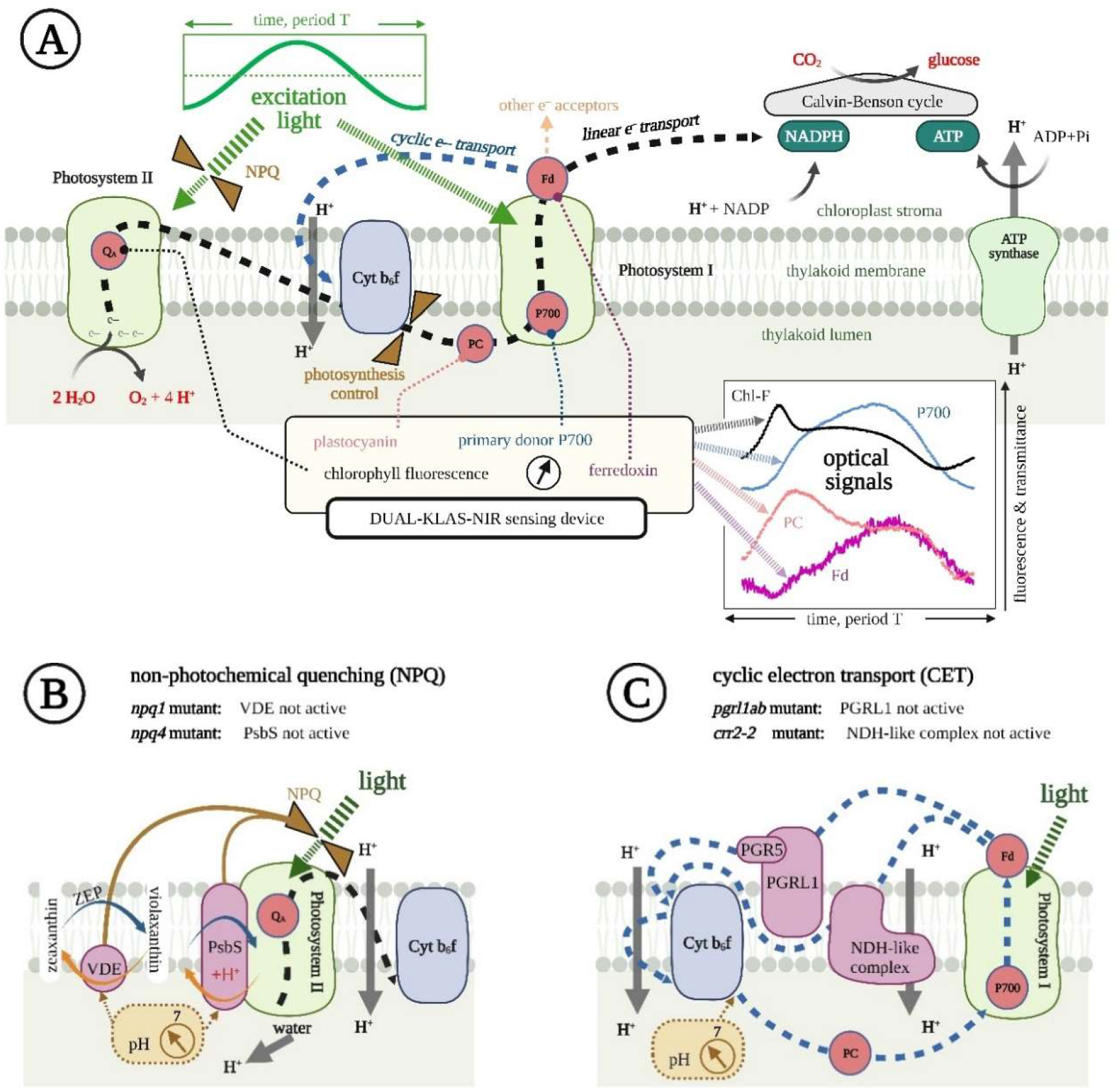
A scheme of the photosynthetic apparatus in *A. thaliana* showing the linear electron transport from water to NADP+ by the black dashed line and the part of the cyclic electron transport around PSI by the blue dashed line. The components that are monitored by the measured optical proxies (bottom right) were: the Q_A_ redox state determining largely the chlorophyll fluorescence yield (ChlF), plastocyanin (PC), primary donor of PSI (P700), and ferredoxin (Fd). The lumen pH-controlled regulation of PSII light harvesting efficiency (NPQ) and of the electron flow from Cyt b6f (photosynthesis control) are indicated by the brown valve symbol. The harmonically modulated light is represented by one period (green line top left). The optical signals measured (bottom right) by the Dual-KLAS-NIR instrument are described in Materials and Methods. **1B.** The violaxanthin de-epoxidase (VDE) and PsbS protein constitute rapid non-photochemical quenching mechanisms that are reducing the excitation of PSII. Both processes are induced by low luminal pH. The cycles with blue- and orange-colored arrows represent transitions between low- and high-light states and back that occur periodically during the light oscillations. Orange arrows indicate NPQ induction processes, triggered by increasing ΔpH, while blue arrows indicate the corresponding NPQ relaxation processes. **1C.** The cyclic electron transport proceeds by two parallel pathways: one *via* the Proton Gradient Regulation 5 (PGR5) and Proton Gradient Regulation-like 1 (PGRL1) complexes and the other *via* the NADH dehydrogenase-like complex (NDH-like) complex. The mutant *pgrl1ab* lacks PGRL1-PGR5-dependent pathway, whereas *crr2-2* lacks the pathway that depends on the NDH-like complex. Both CET pathways lower the luminal pH by electron-coupled proton transport (Shikanai, 2014). In all schemes, the down-pointing gray arrows represent all processes leading to the accumulation of protons in the lumen relative to the stroma. The up-pointing gray arrow shows the dissipation of this potential difference by ATP-synthase.

CET mediates the electron transport from the reduced ferredoxin (Fd) at the acceptor side of PSI *via* plastoquinone pool, Cyt b6f and PC back to the donor side of PSI (Fig.1C). By the electron-coupled proton transport, CET contributes to the proton motive force for ATP synthesis, thus adjusting the ATP/NADPH ratio for downstream carbon assimilation. The CET-induced lumen acidification also contributes to NPQ, and regulates electron transport (Wang *et al.*, 2015), protecting both PSII and PSI (Suorsa *et al.*, 2012; Suorsa *et al.*, 2013; Yamori *et al.*, 2016; Shimakawa & Miyake, 2018; Nakano *et al.*, 2019; Yamamoto & Shikanai, 2019). As in other angiosperms, *A. thaliana* plants possess two CET pathways (Shikanai, 2014). One involves the Proton Gradient Regulation 5 (PGR5) and Proton Gradient Regulation-like 1 (PGRL1) proteins (Munekage *et al.*, 2002; DalCorso *et al.*, 2008; Hertle *et al.*, 2013; Sugimoto *et al.*, 2013) whereas in the other, the electron is transported from Fd to plastoquinone *via* the NADH dehydrogenase-like complex (NDH-like) (Fig.2C) (Yamamoto *et al.*, 2011; Peltier *et al.*, 2016). This latter pathway acidifies the thylakoid lumen not only by the plastoquinone/plastoquinol proton transport, but the NDH-like complex operates also as an energy-coupled proton pump (Kouřil *et al.*, 2014; Strand *et al.*, 2017; Laughlin *et al.*, 2020).

**Fig.2.**
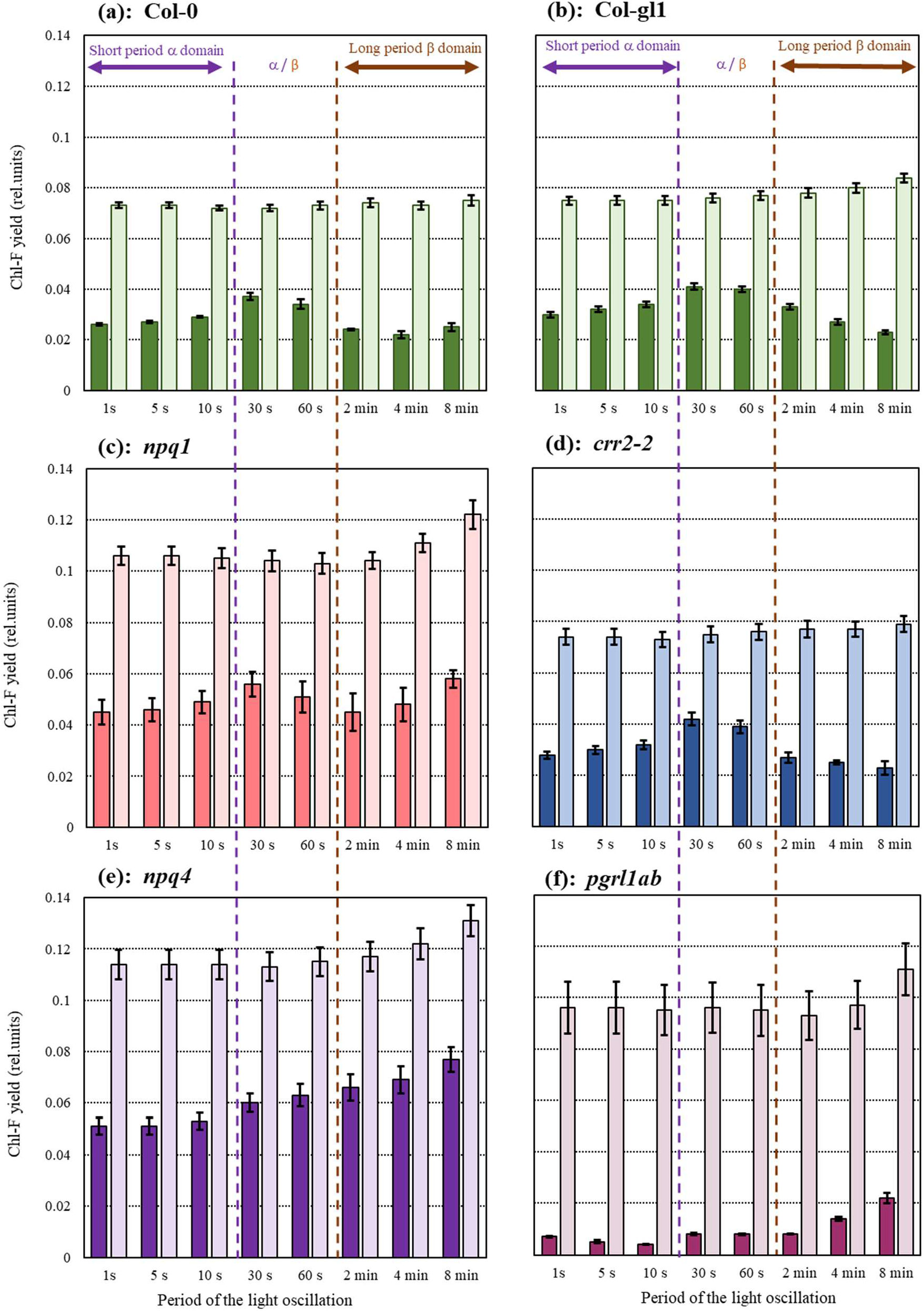
The stationary (light-shaded) and oscillatory (dark-shaded) components of ChlF yield measured in 6 genotypes of *A. thaliana* (n=3): (a) WT Col-0, (b) WT Col-gl1, (c) *npq1,* (d) *crr2-2,* (e) *npq4,* (f) *pgrl1ab.* The actinic light was oscillating with the short periods of 1, 5, and 10 s (α domain), with periods of the 30 and 60 s (boundary α /β domain), and with the long periods 2, 4, and 8 min (β domain). The oscillatory component was calculated as a difference between maximum and minimum of ChlF in one light period. The error bars represent standard errors obtained from three biological replicates.

<u>The first of the objectives</u> of the present study was to identify the limiting frequencies at which the NPQ regulation responds and beyond which it ceases responding, i.e., to determine the NPQ operation range. In contrast to the response times obtained from activation during a dark-to-light induction, e.g., in (Wehner *et al.*, 2004; Nilkens *et al.*, 2010), the limits of the NPQ operation range in oscillating light depend on an interplay of the activation and deactivation processes that both contribute to plant response to fluctuating light. Towards this goal, we explored the responses to oscillating light of the wild-type, the *npq1* mutant, which cannot convert violaxanthin into zeaxanthin by VDE (Niyogi *et al.*, 1998), and of the *npq4* mutant that lacks the PsbS protein (Li *et al.*, 2000).

<u>The second objective</u> was to discriminate between the roles played in an oscillating light by the two parallel pathways of CET. For this, we contrasted the *pgrl1ab* mutant that lacks the (PGRL1/PGR5)-dependent pathway (DalCorso *et al.*, 2008), and the *crr2-2* mutant impaired in the NDH-like complex-dependent pathway (Hashimoto *et al.*, 2003) with their respective wild types.

Understanding of the dynamics in complex systems like plants requires simultaneous measurements of multiple system variables (Ganusov, 2016; Nedbal & Lazár, 2021) that would enable building robust mathematical models in future (Kitano, 2001). <u>Towards this third objective</u>, we studied the dynamics of ChlF, and redox changes in P700, plastocyanin (PC), and ferredoxin (Fd) appearing in plants exposed to light that oscillated between light-limited and saturating intensities with periods changing between 1 s and 8 min. Two frequency domains and the boundary range between them were, in this way, characterized by highly contrasting dynamic behavior of NPQ and CET.

## Materials and Methods

### Plant Material and Growth Conditions

*Arabidopsis thaliana* wild-type Col-0, trichome-lacking wild-type Col-gl1, and the mutants affected in NPQ (*npq1, npq4*) and in CET pathways (*pgrl1ab, crr2-2*) were used in the study. Col-0 was the background of the *npq1, npq4,* and *pgrl1ab* mutants, and the Col-gl1 was the background of *crr2-2.* The mutant *crr2-2* was kindly provided by Toshiharu Shikanai, Kyoto University, Japan and *pgrl1ab* by Dario Leister, Ludwig Maximilian University München, Germany. Seeds were sown in commercial substrate (Pikier, Balster Einheitserdewerk, Fröndenberg, Germany). After 3 days of stratification in a 4°C dark room, the seedlings were transferred to a climate chamber with a light intensity of approx. 100 μmol photons·m^-2^·s^-1^, 12 h/12 h light/dark photoperiod, 26°C/20°C day/night air temperature, and 60% relative air humidity. On the 15^th^ day after sowing, the seedlings were transferred to 330 ml pots, one plant per pot filled with a commercial substrate (Lignostrat Dachgarten extensive, HAWITA, Vechta, Germany). The environmental conditions in the climate chamber remained the same as for the seedlings. The plants were watered every day from the bottom to keep soil moisture approximately constant throughout the cultivation and during the experiments.

### Chlorophyll fluorescence and KLAS-NIR measurements

In the sixth week after sowing, chlorophyll fluorescence yield and redox changes of PC, P700, and Fd were measured simultaneously using a Dual-KLAS-NIR spectrophotometer with a 3010 DUAL leaf cuvette (Heinz Walz GmbH, Effeltrich, Germany). Red actinic light (630 nm) was applied to both sides of the leaf. The pulse-amplitude-modulated green light (540 nm, 6 μmol photons·m^-2^·s^-1^) was applied to the abaxial side of the leaf to excite chlorophyll.

In addition to chlorophyll fluorescence yield, the Dual-KLAS-NIR allows to measure four dual-wavelength difference signals of transmittance simultaneously in the near-infrared part of the spectrum using 780-820, 820-870, 870-965, and 840-965 nm wavelength pairs (Klughammer & Schreiber, 2016; Schreiber & Klughammer, 2016). Four difference signals of transmittance can be deconvoluted into contributions corresponding dominantly to redox changes of ferredoxin (Fd), primary donor of PSI (P700), and plastocyanin (PC). The deconvolution relies on the selective differential transmittance spectra of P700, PC and Fd for the four wavelength pairs, which were determined by the routine called Differential Model Plots (Klughammer & Schreiber, 2016; Schreiber & Klughammer, 2016). The plots were always constructed with 2-3 replicates. The Differential Model Plot was in this way determined for each genotype and used for the respective spectral deconvolution.

Plants were dark acclimated overnight and then remained in darkness until their dynamic properties were investigated by the Dual-KLAS-NIR measurement with constant and oscillating actinic light. The measurement started with the NIR-MAX routine on a dark-acclimated plant to estimate the maximum oxidation of PC and P700 (100%), and maximum reduction of Fd (−100%) as described in (Klughammer & Schreiber, 2016). The deconvolution of the overlapping optical transmission signals in near-infrared and the subsequent quantitative interpretation may, to some extent, be compromised by long measuring times, during which the initial deconvolution and min-max parameters may drift due to changing optical properties of the leaf. This would be of utmost importance in long conventional time-domain measurements that rely on stable reference signals, as time-domain measurements are difficult to correct for drift. With harmonically modulated light, one measures relative changes of the redox states of P700, PC, and Fd that are induced by oscillating light and this method is less sensitive to slowly drifting reference levels. The redox changes are quantified here relative to the value found at the trough of the oscillation, i.e, when the light intensity reaches its minimum. The minimum PAR was always 100 μmol photons·m^-2^·s^-1^. By using a reference level taken in every oscillation period, we reduced or eliminated influence of signal drift that may occur over long experimental periods. The dynamic trends occurring during the oscillations were, in this way, assessed using relative changes of the optical proxies named: apparent relative P700 and PC oxidation, and Fd reduction, respectively.

It is worth noting specific methodological aspects of the Fd signal. The implicit assumption of the NIR-MAX routine is that Fd is fully oxidized in darkness. This may not be always correct, because Fd can be reduced by multiple pathways, even in the absence of light. This potential caveat may result in an incorrect calibration of the optical proxy and in an apparent drift of the optical signal that is ascribed to fully oxidized Fd. In this situation, Fd-reduction may exceed the expected range of 0 to −100%. Therefore, we shall not base any conclusions here on the absolute numerical values of the Fd-redox state but rather focus on trends in the Fd-reduction dynamics relative to the value found at the minimum PAR (100 μmol photons·m^-2^·s^-1^). Further, it is worth noting that when the charge separation induced by light occurs in the PSI core complex, electron transfer from P700 to Fd occurs in a series of rapid redox reactions through A0 (the monomeric form of Chl **a**) and A1 (phylloquinone) to the [4Fe-4S] clusters (FX, FA, and FB), and ultimately to Fd [2Fe-2S] (Schreiber & Klughammer, 2016). In the case of *in vivo* measurements, distinguishing the absorption changes of Fd from that of the other FeS proteins is practically impossible, as the NIR differential spectrum of FA-FB is similar to that of the Fd (Sétif *et al.*, 2019), and much larger absorption changes caused by other components can influence the signal deconvolution of different FeS proteins. Therefore, the “Fd” signal used in this study is a mixture of signals of FeS components at the PSI acceptor side.

### Actinic light protocols

#### Forced oscillations with changing frequencies

Following induction in constant actinic light of 450 μmol photons·m^-2^·s^-1^ that lasted 10 min, the plants were exposed to light that was oscillating around this level, between 100-800 μmol photons·m^-2^·s^-1^. The frequency (periods) and the number of periods of sinusoidal light were set in the KLAS-100 software (Heinz Walz, GmbH, Effeltrich, Germany). The periods were changing in a single continuous sequence: three periods of 8 min, five periods of each 4 min, 2 min, 1 min, 30 s, and 10 s, and finally ten periods of 5 s and 1 s (*Supporting Information Fig. SI-4*). The order in which the periods were changing was not affecting our conclusions and the same results were obtained when the periods were in an increasing sequence (*Supporting Information – Influence of light history on the signal patterns*). Three biological replicates of each *A. thaliana* genotype were examined.

#### Analysis of the signals induced by oscillating light

The first period of 8 mins oscillation was largely influenced by the transition from constant to oscillating light and the first two periods of the other oscillations were influenced by the change of light frequency. Because of these reasons, they were not analyzed (one example is offered in *Supporting Information Fig.SI-4).* The later signals were already periodic and were used to extract the respective dynamic features by numeric analysis. The data representing each respective frequency in multiple replicated plants were averaged to improve the signal-to-noise ratio. The signal averages were then numerically approximated by a function *Fit*(*t*), consisting of a fundamental mode and of 3 upper harmonic modes as described in (Nedbal & Lazár, 2021):

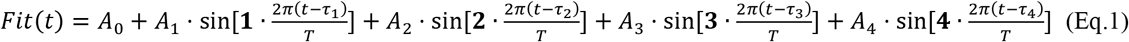

The least-square fitting procedure was done in MS Excel and the fit yielded the stationary component *A*_0_, and the amplitudes and phase shifts {*A*_1_ *τ*_1_} for the fundamental harmonics, {*A*_2_,*τ*_2_}, {*A*_3_,*τ*_3_}, {*A*_4_,*τ*_4_} for the upper harmonic components. Typically, no more than 2 upper harmonics were needed as adding the third upper harmonic mode did not improve χ^2^ of the fit. The fitted signals of P700, PC, and Fd apparent redox changes were normalized by dividing by the signals obtained at the minimum light level (100 μmol photons·m^-2^·s^-1^) as explained above.

## Results

### Chlorophyll fluorescence emission

Eight different oscillation periods of light were sequentially applied to identify the characteristic frequency limits of NPQ, and CET. Plants responded to the light changes by ChlF that was alternating in oscillations between minima and maxima around a stationary level *A*_0_ (Eq. 1). The oscillatory ChlF component was calculated as the difference between the respective minimum and maximum of ChlF during one cycle. The stationary as well as oscillatory ChlF components (Fig.2) were dependent on the period, i.e., frequency of light oscillations in three contrasting ways that can be categorized in the α (frequencies between 1 - 1/10 s^-1^), and β (frequencies between 1/120 - 1/480 s^-1^) domains, and α/β boundary domain, its frequencies 1/30 and 1/60 s^-1^ leading to a distinct resonance in ChlF (*Supporting Information - Introduction*). The categorization of the frequency domains is compatible with that introduced for green algae in (Nedbal & Lazár, 2021).

The domain α included the shortest periods of light oscillations. Although the differences between the genotypes were large, the α domain can be defined by ChlF response remaining for each particular genotype largely the same, no matter if the period was 1, 5, or 10 s (Fig.2). In the boundary α/β domain, the oscillatory ChlF component was typically higher than in the neighboring domains α and β (Fig.2). This local maximum was reported earlier (Nedbal & Březina, 2002) and attributed to a resonance due to a regulatory feedback. This feature, pronounced in WTs, was absent in the *npq4* PsbS-deficient mutant and was damped, probably by electron transport limitation, in the *pgrl1ab* mutant (see arguments below). Both the stationary and oscillatory components of the ChlF yield were increasing in the β domain in the *npq1, npq4,* and *pgrl1ab* mutants when the period increased from 2 min to 8 min (Fig.2). This trend was absent in WTs and *crr2-2.*

The oscillatory components of ChlF in the α, α/β, and β domains are characterized in Fig.2 only by the difference between the minima and maxima. These extremes often appear before or after the minima and maxima of light and Fig.2 does not show the respective phase shift of ChlF. We therefore present additional information about the phase shift of the ChlF extremes and further dynamic details in Fig.3. This figure shows by a color code the entire dynamics of the oscillatory ChlF signal as it develops with the phase of the light oscillation along the abscissa axis and how it depends on the period of the oscillation along the ordinate axis. The ChlF signal was mostly more complex than a simple harmonic variation, containing strong upper harmonics, often rich in phases with ChlF decreasing in increasing light and *vice versa.*

**Fig.3.**
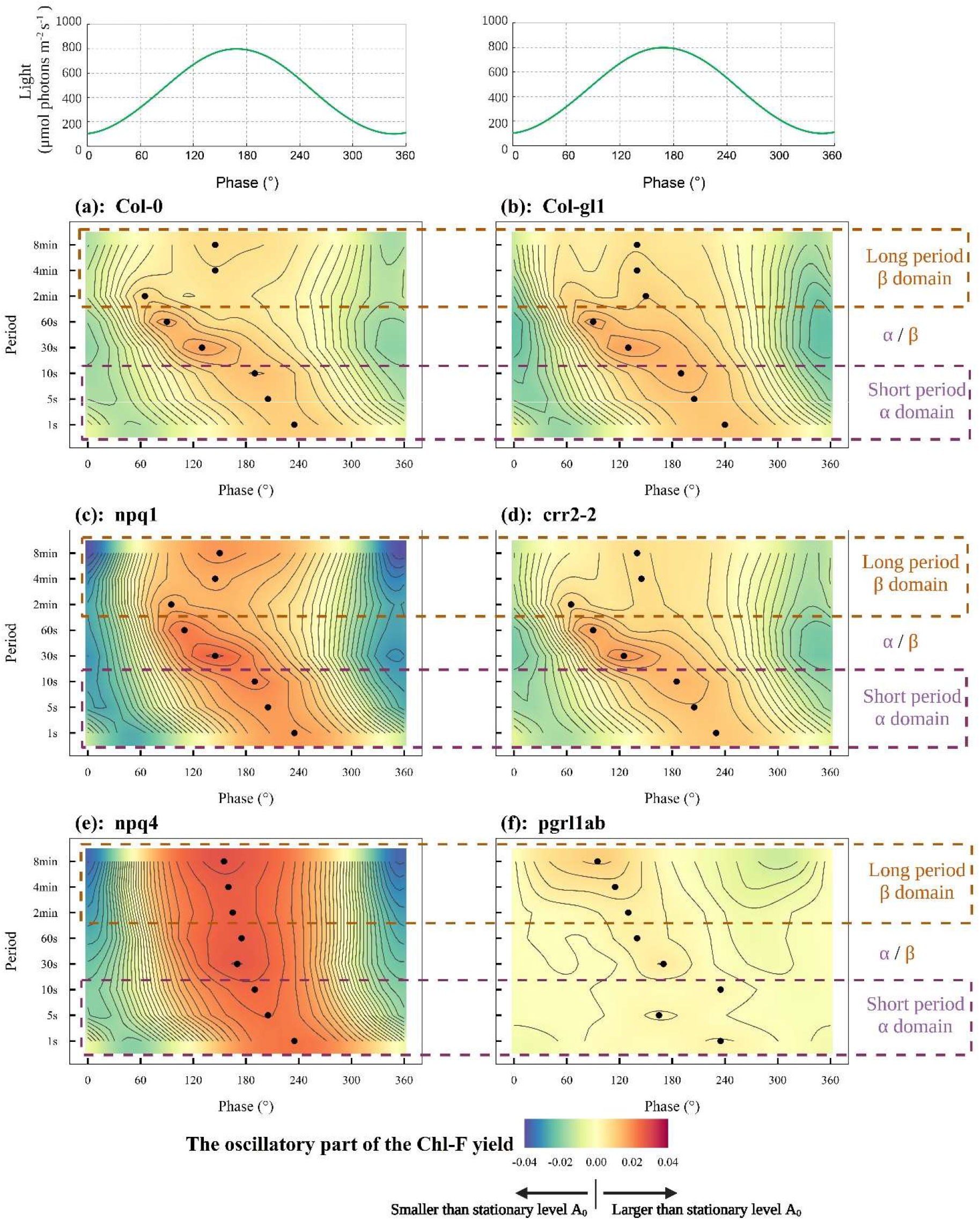
The oscillatory part of the relative chlorophyll fluorescence (ChlF) yield. The green line in the top panel represents light that was oscillating between 100 and 800 μmol photons·m^-2^·s^−1^ with periods in the range of 1 s to 8 min. The progress of the light oscillation and of ChlF response are shown by the phase between 0 and 360°, so that the abscissa is the same for all periods of oscillations that were applied. The other six panels show the oscillatory part of the ChlF yield induced in 6 genotypes of *A. thaliana* (n=3): (a) WT Col-0, (b) WT Col-gl1, (c) *npq1,* (d) *crr2-2,* (e) *npq4,* (f) *pgrl1ab.* Blue colors indicate signals that were below the stationary ChlF yield, representing negative difference of the relative ChlF yield. Red colors represent oscillatory signals above the stationary ChlF yield. The black dots indicate the phase at which maximum ChlF yield at the given period occurred. The contour lines separate signal ranges of 0.0025 of relative units. The brown dashed rectangles indicate long periods of 2, 4 and 8 min (β domain), and the purple dashed rectangles indicate short periods of 1, 5, and 10 s (α domain).

The changes of the ChlF yield in Fig.3 were always induced by the same light pattern (top green line), oscillating between 100 μmol photons·m^-2^·s^-1^ and 800 μmol photons·m^-2^·s^-1^. The line shows the true light pattern that was slightly shifted in phase against the instrument protocol that aimed at generating the light minima at 0° and 360° and maximum at 180°. This instrumental deficiency had no impact on our conclusions.

The data show that both the stationary (Fig.2) as well as the oscillatory part of the ChlF (Figs.2,3) were strongly reduced by NPQ in WT relative to the *npq1* and *npq4* mutants. The most contrasting with WT was the *npq4* PsbS-deficient mutant exhibiting high amplitude ChlF variations (Fig.3). The ChlF followed closely the light intensity in the *npq4* mutant suggesting dominance of photochemical quenching with little interference of NPQ. ChlF was simply increasing due to reducing of the plastoquinone pool in increasing light and decreasing when the pool became more oxidized in decreasing phase of the light oscillation. The amplitude of the ChlF oscillations was monotonously increasing from shorter to longer periods. This observed dynamics of the *npq4* mutant can be explained by the PsbS protein being necessary for rapid NPQ regulation in light that was oscillating with periods as short as 30 s.

Unlike the *npq4* PsbS-deficient mutant, the *npq1* VDE-deficient mutant exhibited a distinct ChlF yield maximum in the α/β boundary. This difference between the *npq* mutants, i.e., the suppression of the ChlF yield maximum in *npq4* in the α/β boundary domain was visible already in Fig.2. Fig.3 shows this difference between *npq4* and other genotypes in further detail. The distinct ChlF maximum in the α/β boundary was found in all biological replicates of all genotypes that were competent in the PsbS-dependent NPQ mechanism and was largely supressed in the *npq4* PsbS-deficient mutant. This PsbS-dependent resonance maximum occurred with the 30 s period in both wild types at 130° phase, i.e., ca. 10.8 s after the light started rising from its minimum and, with the 60 s period, at 90° phase, i.e., ca. 15 s after the light minimum. The phase shift indicates that the plants and their PsbS-dependent NPQ mechanism required 10 - 20 s to change the trend from rising to declining ChlF in the ascending light phase. More on the resonance phenomenon is available in *Supporting Information - Introduction.*

The differences between the *npq1* mutant and WT that were representing dynamic effects of VDE-dependent NPQ in WT were mostly subtle (compare Figs.3a with c). Namely, the local maximum in the α/β boundary domain was present in both. The oscillatory part of the ChlF was in *npq1* much larger than in WT at all frequencies. The same was true for the stationary ChlF (compare Figs.2a with c). The major contrast between the *npq1* mutant and WT was found when the period of the oscillating light increased from 60 s to 2 min and longer (β domain). The oscillatory part of the ChlF yield was largely quenched in the WT while it was rising towards long periods in the VDE-deficient *npq1* mutant. We conclude that the VDE-dependent NPQ was able to respond to light that was oscillating with periods of 2 minutes and longer. In contrast to the dynamic NPQ response, the stationary component of ChlF yield was always quenched in the WT when compared to the *npq1* mutant (Fig.2 and *Supporting Information Fig.SI-5*). We conclude that the VDE-dependent NPQ is largely responsible for protection in static or slowly changing light.

A unique response to oscillating light was found in the *pgrl1ab* mutant that is impaired in the PGR5-PGRL1 pathway. The stationary part of ChlF was suppressed in *pgrl1ab* much less than in WT or in the *crr2-2* mutant (Fig.2, *Supporting Information Fig.SI-5*) while the variations around this stationary level that were caused by the light oscillations were by far smallest among all the tested genotypes (Fig.3). The high stationary and low oscillatory ChlF components can signal congestion of the electron transport chain on the acceptor side of PSII. Unlike in WT and in the *crr2-2* mutant, the variations of ChlF yield in the *pgrl1ab* mutant exposed to long-period light oscillations (β) were stronger than in the short-period oscillations (α) indicating that NPQ became less effective when the periods were long. The dynamic features found in the *pgrl1ab* mutant cannot be attributed solely to changes in CET and it is likely that the lack of the PGRL1 protein impaired also the linear electron transport and affected induction of NPQ and photosynthesis control (DalCorso *et al.*, 2008; Suorsa *et al.*, 2016; Wada *et al.*, 2021).

The ChlF patterns found with the *crr2-2* mutant were like WT (Fig.2,3) suggesting that the response to oscillating light was only marginally affected by knocking out of the NDH-like complex-dependent pathway of CET.

### Transmission proxies of PC, P700, and Fd redox states

The frequency responses of apparent relative oxidation/reduction of P700 (Fig.4), PC (Fig.5), and Fd (Fig.6) in oscillating light revealed diverging dynamics of components operating close to Photosystem I. The P700 oxidation was, in all genotypes except the *pgrl1ab* mutant, closely following the oscillating light (Fig.4). The amplitude of the P700 oscillations was relatively constant for periods shorter than 60 s (α and α/β) but increasing with long periods from 2 to 8 min (β). Compared to other genotypes tested, the *pgrl1ab* mutant showed qualitatively different frequency responses, in which the P700 redox state was largely independent of the oscillating light (Fig.4f). This lack of variability of the P700 redox state signaled slowing or blockage of electron flow on the acceptor side of PSI in the *pgrl1ab* mutant (Shimakawa & Miyake, 2018). This presumed congestion in the linear electron transport may extend back to PSII and also explain the lack of variability of ChlF that was described above.

**Fig.4.**
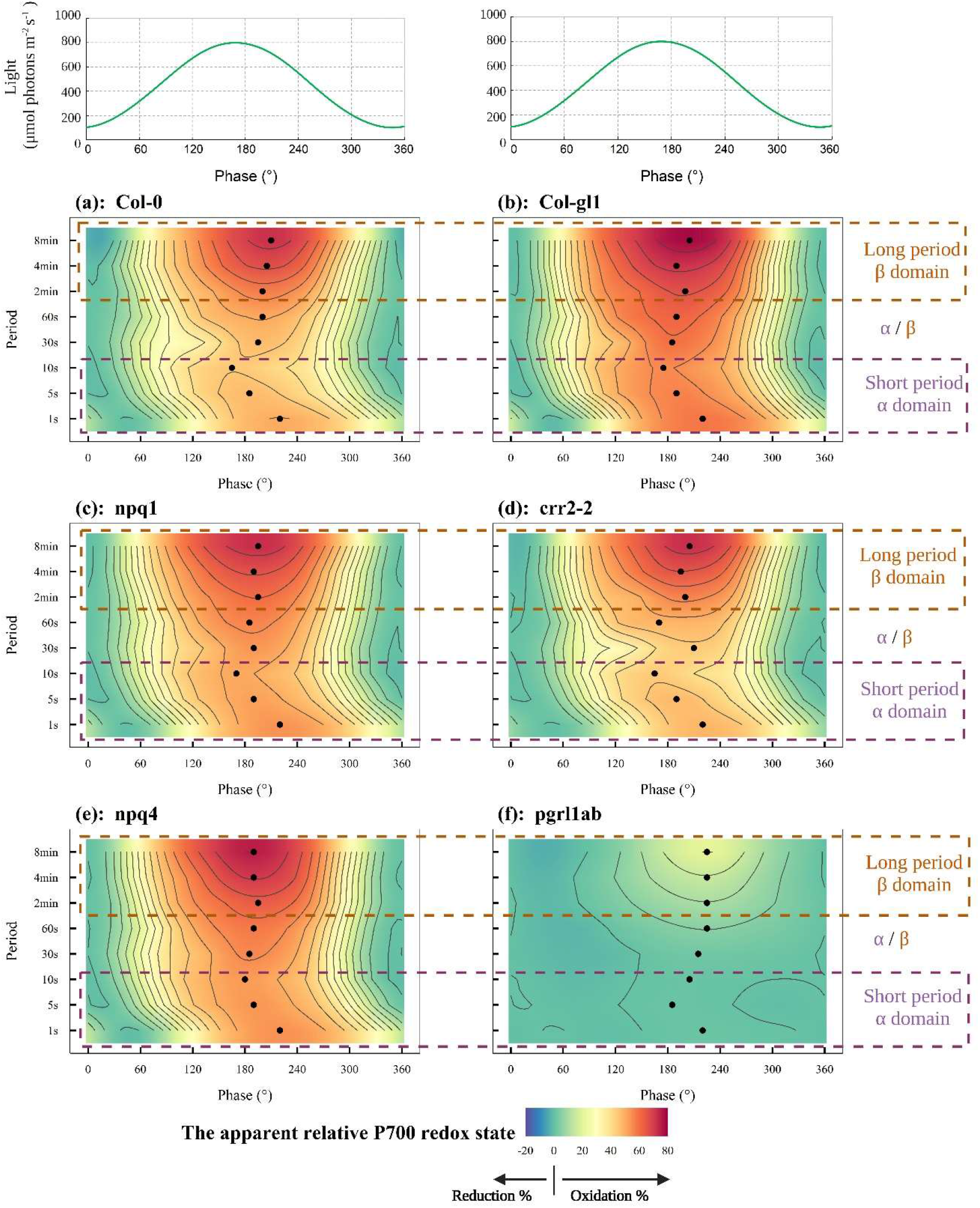
The dynamics of primary donor of PSI (P700) in light that was oscillating as shown by the green line in the top panels. The abscissa shows always the phase of the light oscillation. The six other panels show the apparent changes in the PC redox state in 6 genotypes of *A. thaliana* (n=3): (a) WT Col-0, (b) WT Col-gl1, (c) *npq1,* (d) *crr2-2,* (e) *npq4,* (f) *pgrl1ab*. The dynamics were induced by light oscillating between 100 and 800 μmol photons·m^-2^·s^-1^ with periods in the range of 1 s to 8 min. The black dots indicate at which phase the maximum of the apparent P700 oxidation at a given period occurred. The color scale ranges from blue to red and represents the apparent oxidation of P700 relative to the state at minimum light level 100 μmol photons·m^-2^·^-1^ from low to high. Blue indicates that P700 was more reduced than at the light minimum, while red indicates that P700 was more oxidized than at the light minimum. The brown dashed rectangles indicate long periods of 2, 4 and 8 min (β domain), and the purple dashed rectangles indicate short periods of 1, 5, and 10 s (α domain).

**Fig.5.**
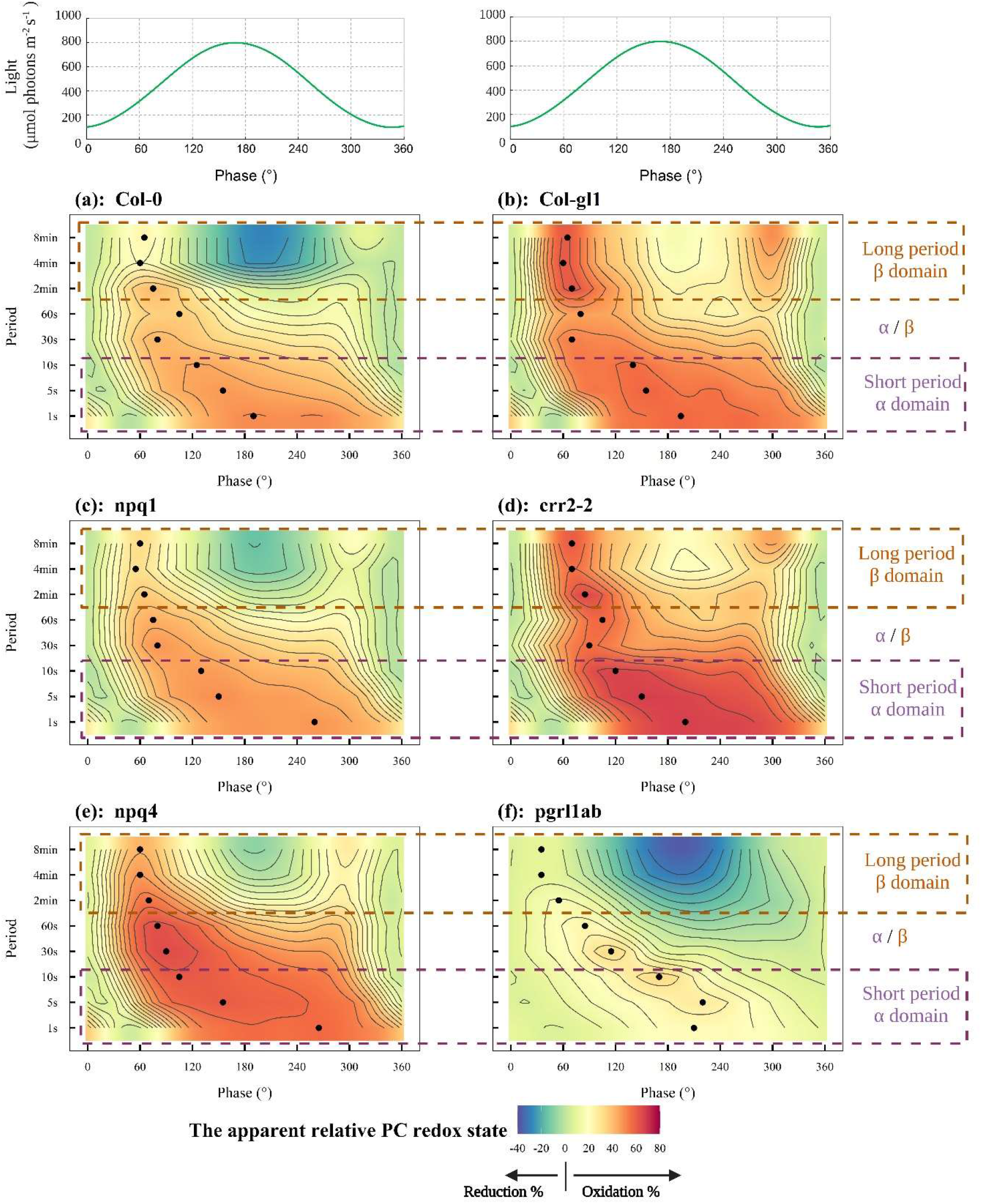
**The plastocyanin (PC) dynamics** in light that was oscillating as shown by the green line in the top panels. The abscissa shows always the phase of the light oscillation. The six other panels show the apparent changes in the PC redox state in 6 genotypes of A. *thaliana* (n=3): (a) WT Col-0, (b) WT Col-gl1, (c) *npq1,* (d) *crr2-2,* (e) *npq4,* (f)*pgrl1ab.* The dynamics were induced by light oscillating between 100 and 800 μmol photons·m^-2^·s^−1^ with periods in the range of 1 s to 8 min. The black dots indicate at which phase the maximum of the apparent PC oxidation at a given period occurred. The color scale ranges from blue to red and represents the apparent oxidation of PC relative to the state at minimum light level 100 μmol photons·m^-2^·s^-1^ from low to high. Blue indicates that PC was more reduced than at the light minimum, while red indicates that PC was more oxidized than at the light minimum. The brown dashed rectangles indicate long periods of 2, 4 and 8 min (β domain), and the purple dashed rectangles indicate short periods of 1, 5, and 10 s (α domain).

**Fig.6.**
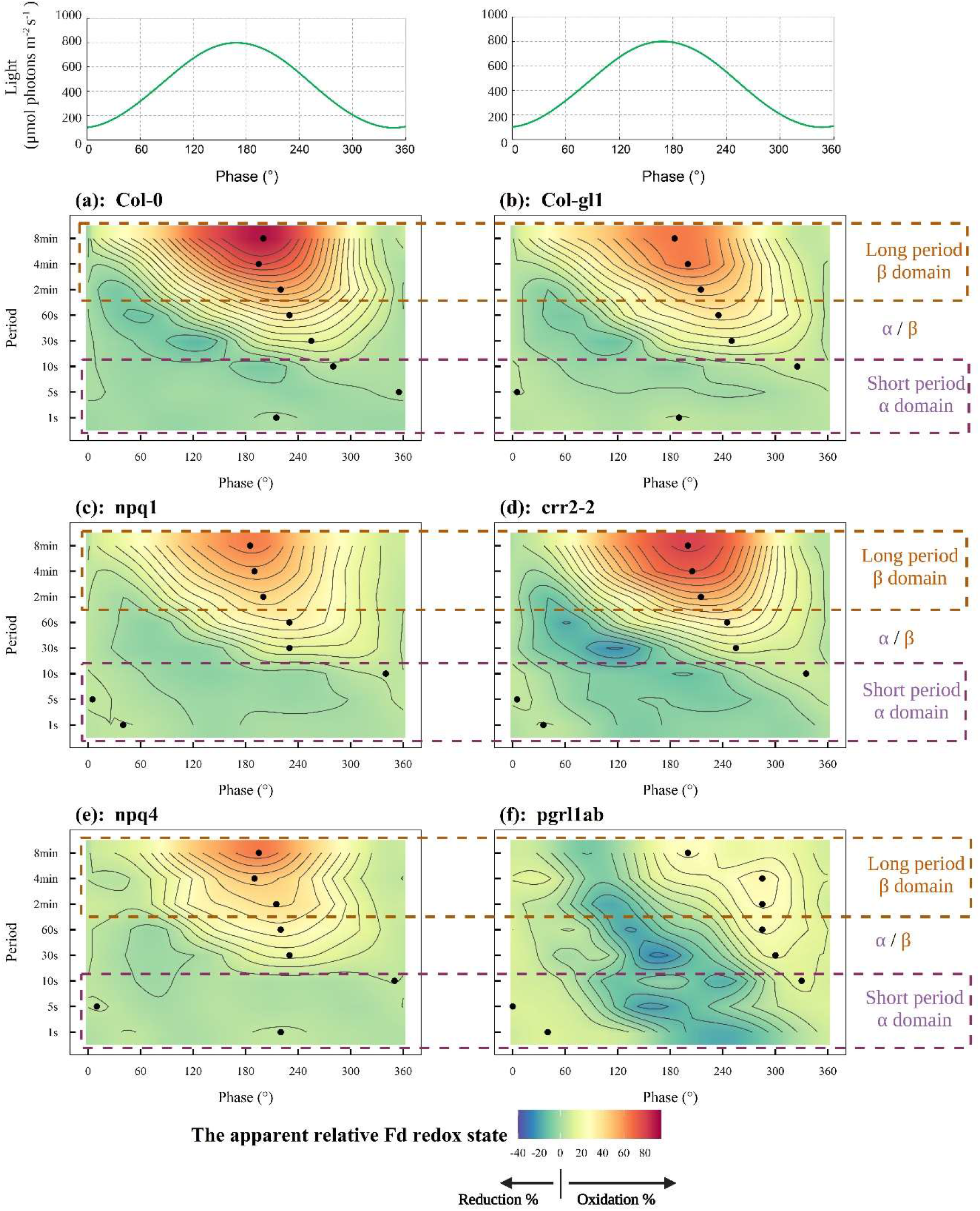
**The ferredoxin (Fd) dynamics** in light that was oscillating as shown by the green line in the top panels. The six other panels show the apparent changes in the Fd redox state in 6 genotypes of *A. thaliana* (n=3): (a) WT Col-0, (b) WT Col-gl1, (c) *npq1,* (d) *crr2-2,* (e) *npq4,* (f) *pgrl1ab.* The dynamics were induced by light oscillating between 100 and 800 μmol photons·m^-2^·^-1^ with periods in the range of 1 s to 8 min. The black dots indicate at which phase the maximum of the apparent Fd oxidation at a given period occurred. The color scale ranges from blue to red and represents the apparent oxidation of Fd relative to the state at minimum light level 100 μmol photons·m^-2^·s^-1^ from low to high. Blue indicates that Fd was more reduced than at the light minimum, while red indicates that Fd was more oxidized than at the light minimum. The brown dashed rectangles indicate long periods of 2, 4 and 8 min (β domain), and the purple dashed rectangles indicate short periods of 1, 5, and 10 s (α domain).

The frequency responses of the apparent relative PC oxidation (Fig.5) depended strongly on the oscillation periods in all genotypes. In a rapidly oscillating light (α), PC was more oxidized in the high light phase than around the light minima. Slow light oscillations (β) elicited a distinct phase dependence, in which PC was increasingly oxidized only in the first phase of the rising light. This trend was changed at a later ascending phase of the oscillation, when the PC oxidation dropped despite of light still increasing and a saddle-type depression occurred around the light maximum.

The apparent relative Fd redox states (Fig.6) were, in WT and the *npq1* and *npq4* mutants, hardly changing between the light minima and light maxima of the rapid light oscillations (α). The NPQ limitations in the *npq1* and *npq4* mutants had only a minor effect on the redox state dynamics of the PSI primary donor P700 and on the PC and Fd dynamics. In the CET mutants, particularly in the *pgrl1ab* mutant, the apparent relative Fd proxy was signaling increasing reduction on the acceptor side of PSI in the strong light relative to the light minima. An apparent Fd reduction was occurring in the *crr2-2* mutant with the periods of 30 and 60 s, i.e., in the α/β boundary (Fig.6d), where the ChlF yield exhibited a resonance feature in all PsbS-competent genotypes (Fig.3) including *crr2-2.* In the slow light oscillations (β), the apparent relative Fd-proxy signaled a high oxidation on the acceptor side of PSI around the light maxima relative to light minima in all genotypes except the *pgrl1ab* mutant.

## Discussion

The regulation of plant photosynthesis has its operating frequency range in which the contributing molecular mechanisms can function. This study employed harmonically oscillating light with different frequencies to identify the frequency limits of multiple photosynthetic regulatory processes. When the light oscillations exceed these limits being, e.g., faster than the limit, the regulation responds only to the average of the light but cannot compensate for the rapidly changing light intensity. The transition at the limiting frequency of the operating range was manifested here by the changing dynamic response of plant photosynthesis, here ChlF, and the PC, P700, and Fd proxies, to light oscillations.

This work identified three types of plant responses that occurred in three frequency domains of light oscillations (α, α/β, β) that often occur in plant canopies. The results obtained with the *npq1* and *npq4* mutants led to formulating the hypotheses on two frequency limits of PsbS- and VDE-dependent NPQ that limit the operation range in natural fluctuating light: the first separating the domain α from α/β at about 1/10 - 1/30 s^-1^ and the second between the domains α/β and β, by the separating limit at about 1/60 - 1/120 s^-1^ (Fig.7). The contrasting responses of the *crr2-2* and *pgrl1ab* CET mutants also supported the hypothesis that parallel pathways or mechanisms acting towards the same function respond differently in the α, α/β, and β domains of the oscillating light.

**Fig.7.**
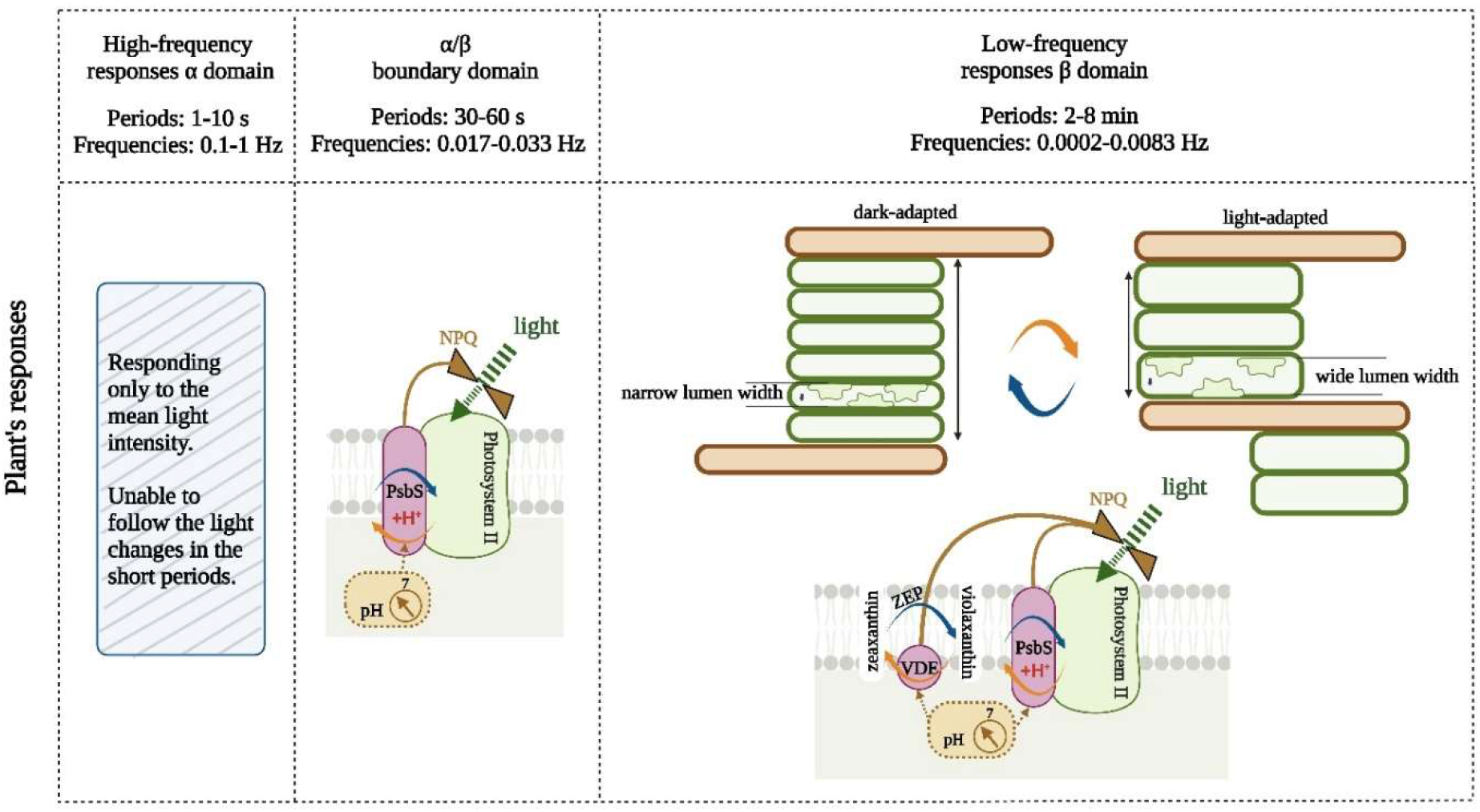
The elemental harmonic components contributing to the natural fluctuations are classified into three frequency domains in which distinct dynamic responses occur (α, α/β, β). The scheme here represents a hypothesis that is proposed to explain the observed frequency responses of WT and *npq1* and *npq4* genotypes in these three domains. The NPQ response in the α/β boundary domain is proposed to be PsbS-dependent. The response to slow oscillations in the β-domain is, in addition formed also by VDE-dependent NPQ that is tentatively proposed to occur with periodic reorganization of the thylakoid membrane. The dark- or low-light adapted thylakoids are organized in large grana with narrow lumen space, reducing the area of contact between the grana membranes (green) and stromal lamellae (brown). With increasing the light intensity towards the maximum of the oscillation in our study, the thylakoids were proposed (Wood *et al.*, 2018; Johnson & Wientjes, 2020) to be reorganized into numerous smaller grana, creating a larger area of contact between the grana membranes (green) and stromal lamellae (brown). Thylakoid lumen volume is proposed to expand with increasing light intensity and shrink when the light is decreasing (Kirchhoff *et al.*, 2011). The symbols used to describe the processes in the thylakoid membrane are the same as in Fig. 1. The lumen pH-controlled regulation of PSII light harvesting efficiency (NPQ) is indicated by the brown valve symbol. The cycles with orange- and blue-colored arrows represent transitions between low- and high-light states and back that may occur periodically during the light oscillations. Orange arrows indicate NPQ induction processes, triggered by increasing ΔpH, while blue arrows indicate the corresponding NPQ relaxation processes.

The light oscillations in the α domain were rapid and NPQ, being unable to follow the fast changes, responded mainly to the mean irradiance. This reduced both stationary as well as oscillatory components of ChlF. NPQ was strongest in the genotypes that were competent both in the PsbS- as well as VDE-dependent quenching and weakest in the *npq4* PsbS-deficient mutant. The PC, P700, and Fd proxies were also following the rapid light modulation in the α domain with a small delay which can be expected for the unregulated system of reactant pools that are filled and emptied by photosynthetic light reactions (Figs.4,5,6). The delay was larger for the faster light oscillations indicating inertia of the pools. The only exception was the *pgrl1ab* mutant in which the electron transport was presumably congested, eliminating largely any variability (Shimakawa & Miyake, 2018).

The boundary α/β domain, represented here by the periods 30 and 60 s, was characterized by a crest maximum of ChlF that was found in plants competent in the PsbS-dependent NPQ and was absent in the *npq4* mutant missing PsbS (Li *et al.*, 2000). This dynamic feature can be assigned to the onset of the PsbS-dependent NPQ in light periods as short as 30 s. We propose that with these short periods, the VDE-dependent NPQ was unable to respond fast enough and the PsbS-dependent NPQ defined the plant’s dynamic response in this domain alone (Fig.7). The PsbS-dependent regulation activated in the α/β domain, marked by an increase of ChlF may thus be responsible for the resonance identified earlier (Nedbal & Březina, 2002) and also may play a role in the spontaneous oscillations in plants that also occur with a period around 60 s (Delieu & Walker, 1983; Lazár *et al.*, 2005), i.e. within the α/β domain. More on the resonance and spontaneous oscillations is available in *Supporting Information - Introduction*. Despite the striking similarity between the resonance frequency and the frequency of spontaneous oscillations, involvement of PsbS-dependent regulation in the spontaneous oscillations of plants remains to be tested by future direct experiments that would also clarify if there were a synergy with activation of the Calvin-Benson cycle (Kaiser *et al.*, 2016; Graham *et al.*, 2017).

The light oscillations in the β domain were slow and the oscillatory component of the ChlF yield was, in WT, strongly suppressed by PsbS- as well as VDE-dependent NPQ. The redox changes near PSI that were induced by slowly oscillating light occurred in WT with high amplitudes despite efficient NPQ. We propose that the contrast between the PC and P700 redox changes during the slow light oscillations (Figs.4,5) might be caused by a periodic light-induced reorganization of the thylakoids that may differently affect PC and P700 (Ruban & Johnson, 2015; Ünnep *et al.*, 2017; Johnson & Wientjes, 2020; Li *et al.*, 2020; Hepworth *et al.*, 2021). Other alternative explanations for the observed slow changes could be the activation and de-activation of the Calvin-Benson cycle (Kaiser *et al.*, 2016; Graham *et al.*, 2017), malate valve (Thormählen *et al.*, 2017), chloroplast movements (Kihara *et al.*, 2020), or changing stomata conductance (Matthews *et al.*, 2018; Li *et al.*, 2021). The last two alternative mechanisms are however unlikely because we have not used blue light excitation that is known to elicit chloroplast movements and the light changes were in our experiments too fast to be followed by the stomata aperture. Based on our results, slow light oscillations up to the period of 8 min are proposed to elicit periodic changes of PsbS-and VDE-dependent NPQ together with periodically changing thylakoid organization (Fig.7). The effect of such a concerted regulation is small in the PsbS-impaired *npq4* mutant and limited in the VDE-impaired *npq1* mutant, indicating that PsbS, VDE, and thylakoid reorganization (Horton & Ruban, 2005) are required for fully functional regulation in the slowly oscillating light.

According to this model (Fig.7), the PsbS-dependent NPQ plays an indispensable role in the plant’s response to light that oscillates in the α/β and β domains. The *npq4* mutant impaired in this function will, in a natural fluctuating light with a range covering 30 s to 8 min, experience largely undamped oscillations in its electron transport chain and, most likely suffer from a dynamic load stress even though PAR may remain in the oscillations far below high levels that would cause photoinhibition in a static light.

It is important to note that the two frequency limits that were identified here to separate domains with different dynamic behaviour of photosynthetic components (Fig.7) are not equivalent to the rate constants that were earlier identified, e.g., by dark-to-light induction experiments. These rate constants or their light-acclimated values co-determine the frequency limits. The rate constant of antheraxanthin accumulation 1/130 – 1/220 s^-1^ reported in (Wehner *et al.*, 2004) is not far from the lower frequency limit that we found separating the α/β domain from β (1/60 - 1/120 s^-1^). Other work (Nilkens *et al.*, 2010) characterized the NPQ response by two characteristic time constants. In WT, they found a fast component with 70% relative contribution responding with a characteristic time of 60 s and a slower 30% component with 570 s. In mutant plants, the components were 50% and 50% and appeared with characteristic times of 460 and 2600 s in *npq4* and 9 and 1600 s in *npq1.*

One can further illustrate the difference between the rate constants measured in dark-to-light induction experiments and the frequency limits identified here using the VDE-dependent NPQ. The conventional time-domain induction experiments reveal the de-epoxidation of violaxanthin into zeaxanthin whereas the frequency limit found in the oscillating light would depend both on the de-epoxidation in rising light and epoxidation in the declining light. Considering that the epoxidation is typically a slower process (Demmig *et al.*, 1987) than even our longest oscillations, we conclude that the oscillating VDE-dependent NPQ in our experiments does not reflect oscillating zeaxanthin levels. This could change in higher temperatures at or above 25°C when the epoxidation is faster (von Bismarck *et al.*, 2022). We propose as a hypothesis that zeaxanthin was produced largely during the pre-illumination and early phases of the oscillating light but remained at a stationary level without oscillating because the zeaxanthin epoxidation expected when the light was declining was not fast enough to occur within the period. If confirmed, this model would lead to the conclusion that the VDE-dependent oscillatory NPQ required zeaxanthin to be present, but the respective NPQ periodic changes were caused by another agent, not by oscillating zeaxanthin concentrations.

Experiments using the KLAS-NIR spectrometer also revealed that the redox changes in and near PSI that were induced by oscillating light remained highly dynamic despite effective NPQ regulation in WT plants. The dynamic changes around PSI were largely independent of the mutations affecting NPQ. In the slow oscillation β domain, the dynamics of P700 were profoundly different from those of PC (Figs.4,5). This may seem as contradicting the close interaction between the two redox components, with PC shuttling electrons from Cyt b6f to P700. We propose that unlike P700 dynamics, the PC dynamics is modulated by a factor that connects both photosystems, such as photosynthesis control (Suorsa *et al.*, 2013; Colombo *et al.*, 2016; Johnson & Berry, 2021) and/or systemic properties such as changing thylakoid topology (Ünnep *et al.*, 2017; Johnson & Wientjes, 2020; Li *et al.*, 2020; Hepworth *et al.*, 2021) that affects PC diffusion (Kirchhoff *et al.*, 2011). The remodeling of the thylakoid membrane structure in slowly oscillating light may affect pronouncedly the heterogeneity of the plastoquinone pool and Cyt b6f in granal and stromal segments of the thylakoid membranes (Joliot *et al.*, 1992; Kirchhoff *et al.*, 2000; Malone *et al.*, 2021). The PQ pool in the grana thylakoid domains participates in the linear electron transport from PSII to Cyt b6f whereas the PQ pool in the stromal thylakoid domains serves primarily for CET (see Fig. 1). Further, one ought to consider that PC transports electrons from Cyt b6f to P700 over long distance (Höhner *et al.*, 2020), presumably with a low probability of backward electron transfer from P700 to PC and is subject to photosynthesis control (Johnson & Berry, 2021). The proposed effect of the thylakoid remodeling on the photosynthetic dynamics needs to be further examined because it may entail fundamental consequences for studies that are based on the dark-to-light transitions in the time domain. Unlike these conventional experimental protocols, the frequency-domain experiments employing oscillating light are applied to light-acclimated plants rather than to plants that change from dark-acclimated to light-acclimated state.

Identifying and quantifying frequency limits for operation of regulatory mechanisms and alternative pathways in plants exposed to oscillating light holds promise for improvements of photosynthetic efficiency in nature under fluctuating light. This may parallel the tremendous success of applying frequency domain analysis in engineering and other fields of science (Ogata, 2010).

## Acknowledgements

Y.N., S.D.S., and L.N. gratefully acknowledge the financial support from the Federal Ministry of Education and Research of Germany (BMBF) in the framework of the YESPVNIGBEN project (03SF0576A).

D.L. and L.N. were supported by the European Regional Development Fund project “Plants as a tool for sustainable global development” (CZ.02.1.01/0.0/0.0/16_019/0000827).

D.L., S.M., and L.N. gratefully acknowledge the financial support by HORIZON-EIC-2021-PATHFINDER OPEN project DREAM (grant agreement No 101046451)

Work by DMK was supported by the U.S. Department of Energy (DOE), Office of Science, Basic Energy Sciences (BES) under Award no. DE-SC0007101.

## Author Contribution

YN, SM and LN planned and designed the research. YN performed experiments. YN, SDS, and LN analyzed data. YN, DL, ARH, DMK, SM, SDS, and LN interpreted the results. YN, DL, ARH, DMK, SM, FF, HP, SDS, and LN wrote the manuscript.

## Data Availability

Data are publicly available in the manuscript, and in the Supporting Information. Further details are provided in https://doi.org/10.1101/2022.02.09.479783.

## Abbreviations and symbols

| Abbreviations and symbols |  |
| --- | --- |
| Parameter | Description |
| CET | Cyclic electron transport |
| Chl | Chlorophyll |
| ChlF | Chlorophyll fluorescence |
| Cyt b<sub>6</sub>f | Cytochrome b<sub>6</sub>f complex |
| Fd | Ferredoxin |
| NIR | Near-infrared light |
| NPQ | Non-photochemical quenching. Multiple molecular mechanisms lead to the lowering of emission of Chl fluorescence in strong light. Here we use the term referring to mechanisms that are reversible on the scale of seconds to minutes and that involve PsbS and/or VDE proteins |
| NDH-like | Chloroplast NAD(P)H dehydrogenase-like complex |
| P700 | Primary electron donor in the reaction center of photosystem I |
| PAR | Photosynthetically active radiation between 400 and 700 nm, measured in units $\mu\text{mol photons}\cdot\text{m}^{-2}\cdot\text{s}^{-1}$ |
| PC | Plastocyanin |
| PSI | Photosystem I |
| PSII | Photosystem II |
| PGR5 | Proton Gradient Regulation 5 protein |
| PGRL1 | Proton Gradient Regulation-like 1 protein |
| PsbS | Chloroplastic 22 kDa Photosystem II protein involved in NPQ |
| VDE | Violaxanthin de-epoxidase enzyme |
| $\alpha$ frequency domain | The domain with periods shorter than 30 s was identified by a specific dynamic behavior of ChlF |
| $\beta$ frequency domain | Another domain of distinct dynamics with periods longer than 1 min and shorter than 8 min |

## Supporting Information

### SI Introduction: Frequency-domain analysis

Plants grow in light that often fluctuates. The frequencies of the light fluctuations can be categorized in domains in which plants respond similarly. The concept of frequency domains was introduced for green algae in (Nedbal & Lazár, 2021). Below it is schematically represented in *Fig.SI-1* for the three domains investigated in higher plant in this study.

**Fig.SI-1.**
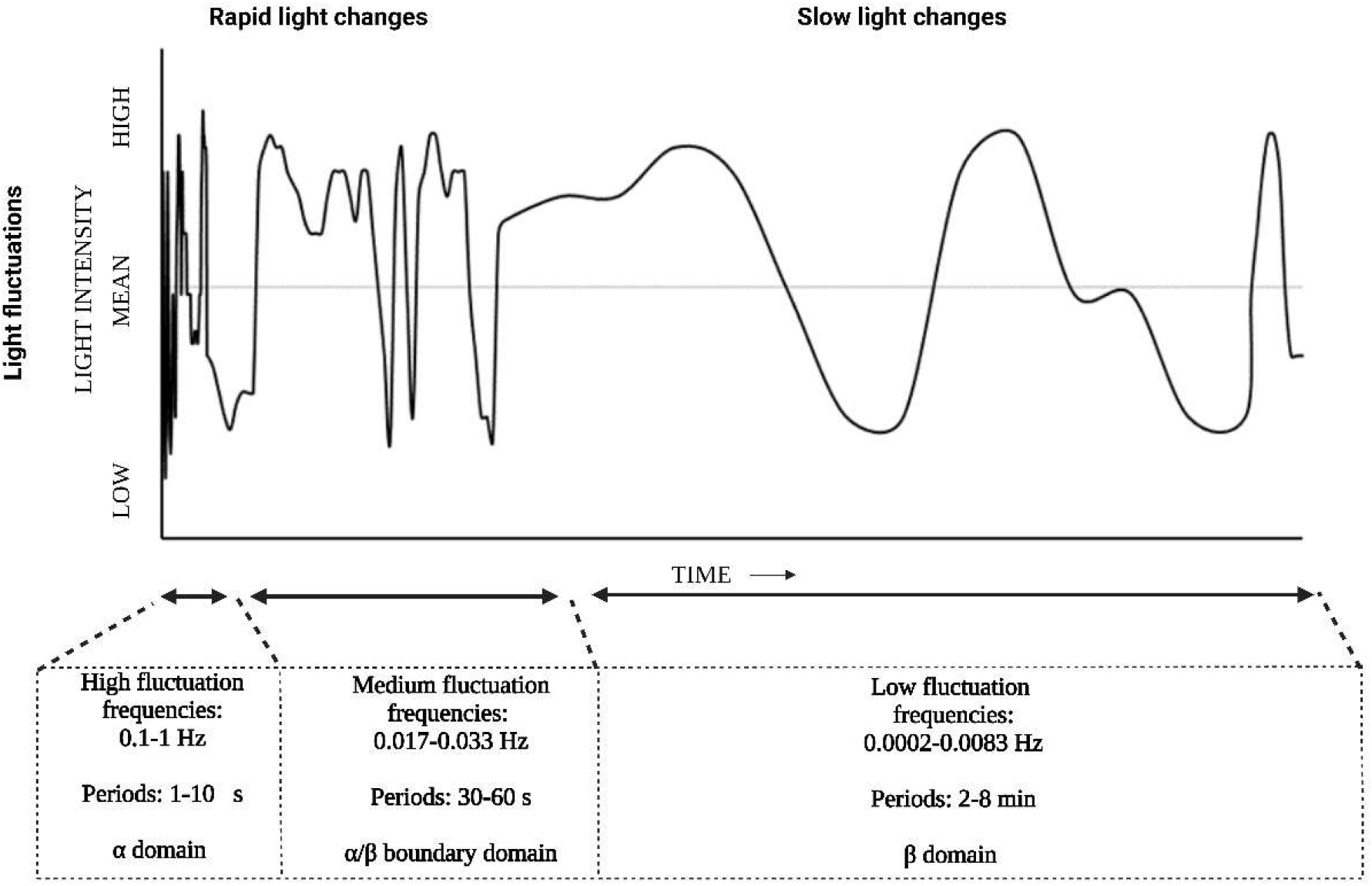
Schematic representation of natural light fluctuations occurring within plant canopies, often caused by transient gaps in the upper canopy, wind induced canopy movement and intermittent cloudiness. The elemental harmonic components contributing to the natural fluctuations are classified into three frequency domains in which distinct dynamic responses occur using nomenclature introduced in (Nedbal & Lazár, 2021).

Any random fluctuation of light can be reproduced as a superposition of harmonic functions, each determined by the period and amplitude of a sinus and cosinus oscillations (Schwartz, 2008). *Fig.SI-2* shows an example of a square sun fleck, e.g., in a forest canopy, that is approximated by seven harmonic components.

**Fig.SI-2.**
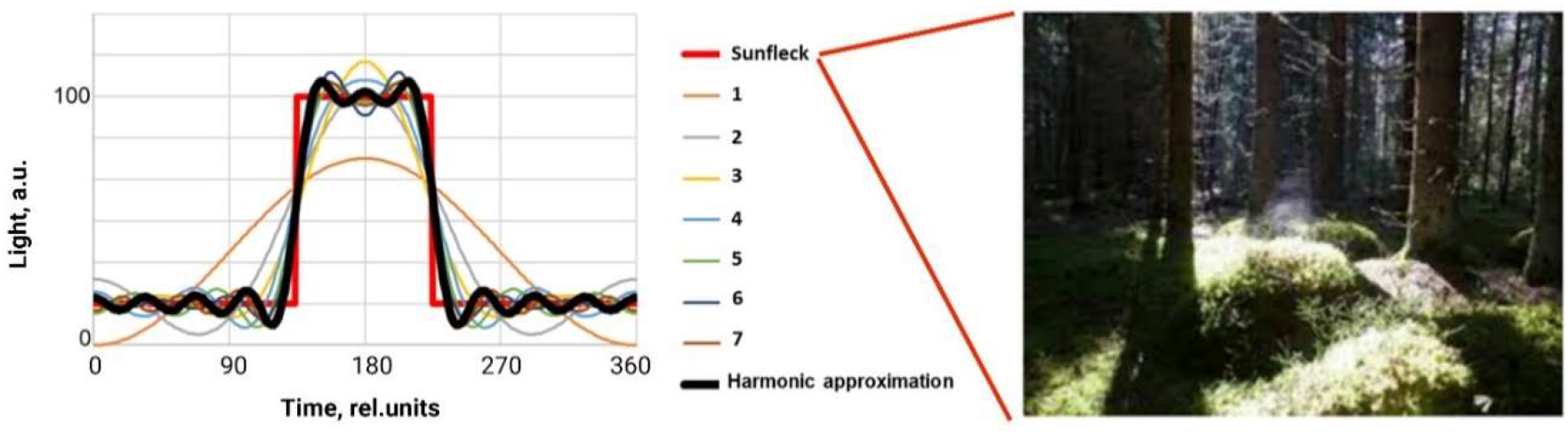
An approximation (thick black line) of a square sun fleck (thick red line) by seven harmonic components. The thin colored lines show how the approximation is improving when the number of contributing harmonic component increases from one (orange, 1^st^ -order) to seven (dark brown, 7^th^ -order).

Natural light fluctuations (e.g., *Fig.SI-1*) do not have perfectly sharp edges and the number of contributing harmonic components would be therefore lower than in *Fig.SI-2.* Once a natural fluctuating light is characterized by characteristic frequencies, one can ask if the photosynthetic regulation can respond to all of them or if some frequencies are outside the operating range of the regulation and cannot be followed by its response. This can be probed in the laboratory by exposing plants to harmonically modulated light of a particular frequency and amplitude. We performed such frequency-domain analysis in the present manuscript for periods between 1 s and 8 min, i.e., for frequencies between 1 s^-1^ and 1/(8·60) s^-1^.

The rate of photosynthesis is linearly proportional to the light intensity only until the constituent redox reactions are saturated. This means, that small increases in light intensity would result in nearly linear increases in photosynthetic electron transfer. If this linear relationship would hold not only for the constant but also for an oscillating light, one could estimate photosynthetic response to an arbitrary light fluctuation by sum of photosynthetic responses to characteristic harmonic components that form the fluctuation. This would mean that the photosynthetic activity occurring during the light fleck in *Fig.SI-2* would be modelled by adding photosynthetic response to the elemental light oscillations that are shown by the thin lines. Knowing photosynthetic reactions for all frequencies appearing in nature would then mean a possibility to predict photosynthetic dynamics in any light pattern that can occur in natural canopy.

This linear approach is limited to very small variations of light intensity. Larger increases in light result in non-linearity for two main reasons. First, downstream reactions tend to become saturated at high light, leading to the well-known light saturation, or “I-P” curves. The second type of non-linearity is caused by downstream processes, especially NPQ and photosynthesis control, or by the onset of metabolic limitations. This is in focus of the present paper as the responses to oscillating light are affected by non-linear processes of NPQ and CET and the limits of their operation frequency range can be identified by scanning through frequencies with light oscillations and identifying frequencies at which the dynamics in WT and relevant mutants abruptly change.

#### Connection between forced and spontaneous oscillations and the resonance phenomenon

Some systems, including plants, oscillate spontaneously even without a periodic stimulation. This phenomenon is sometimes also called autonomous oscillation or self-oscillation. It is the generation and maintenance of a (semi)periodic motion by a source of power that lacks any corresponding periodicity. A very simple example is the classical pendulum. Applying an initial disturbing force results in an extended oscillation behavior with a characteristic natural frequency, sometimes called eigenfrequency. The amplitude of the oscillation damps over time but the natural frequency remains almost constant. This also happens in plants when the air CO2 concentration or light intensity abruptly changes (e.g., in Ferimazova *et al.*, 2002). The plants then start to oscillate with a period of ca. 60 s and the oscillations are damped over ca. 10-15 minutes.

The analogy with a pendulum shows an important relation between the spontaneous oscillations that fade away and the oscillations that are forced and sustained by periodic stimulation. If the pendulum is a part of a clock, there is a mechanism that forces the oscillation to persist over time. This mechanism stimulates the oscillations exactly at the natural frequency of the pendulum. The stimulation would be less effective if the forcing frequency were slower or faster than the natural frequency of the pendulum. The agreement between the natural frequency and the forcing frequency is called resonance. The amplitude of the forced oscillations is always highest when the forcing frequency is in resonance.

Knowing that plants exhibit spontaneous oscillations with a period of around 60 s, one can expect that oscillations forced by oscillating light of the same period will increase the amplitude of, e.g., ChlF. This was exactly the case shown in Figs.2 and 3.

#### Experimental simulation of light fluctuation by square and harmonic modulation

The response of plants to fluctuating light is often experimentally investigated in light that is square-modulated. *Fig.SI-3* shows by red line light that is square-modulated with a period consisting of 1.5 min high light and 1.5 min low light. The light can be approximated by a sum of four harmonics that are shown in *Fig.SI-3* by the thin colored lines. The approximation is shown by the dashed black line. The component with the fundamental harmonic period 3 min is contributing most. The second most contributing is the first upper harmonic component with the period 1 min. The amplitude of the first upper harmonic component (1 min) is three times lower than that of the fundamental harmonic (3 min) but its influence on plant photosynthesis can be enhanced by the resonance described above that occurs for periods around 1 min. With this, the experiments aimed at fluctuating light periodicity of 3 min may well involve an amplified response of the investigated plant to 1 min component. It is not the case when the harmonically modulated light regimes with the periods 3 and 1 min are applied separately.

**Fig.SI-3.**
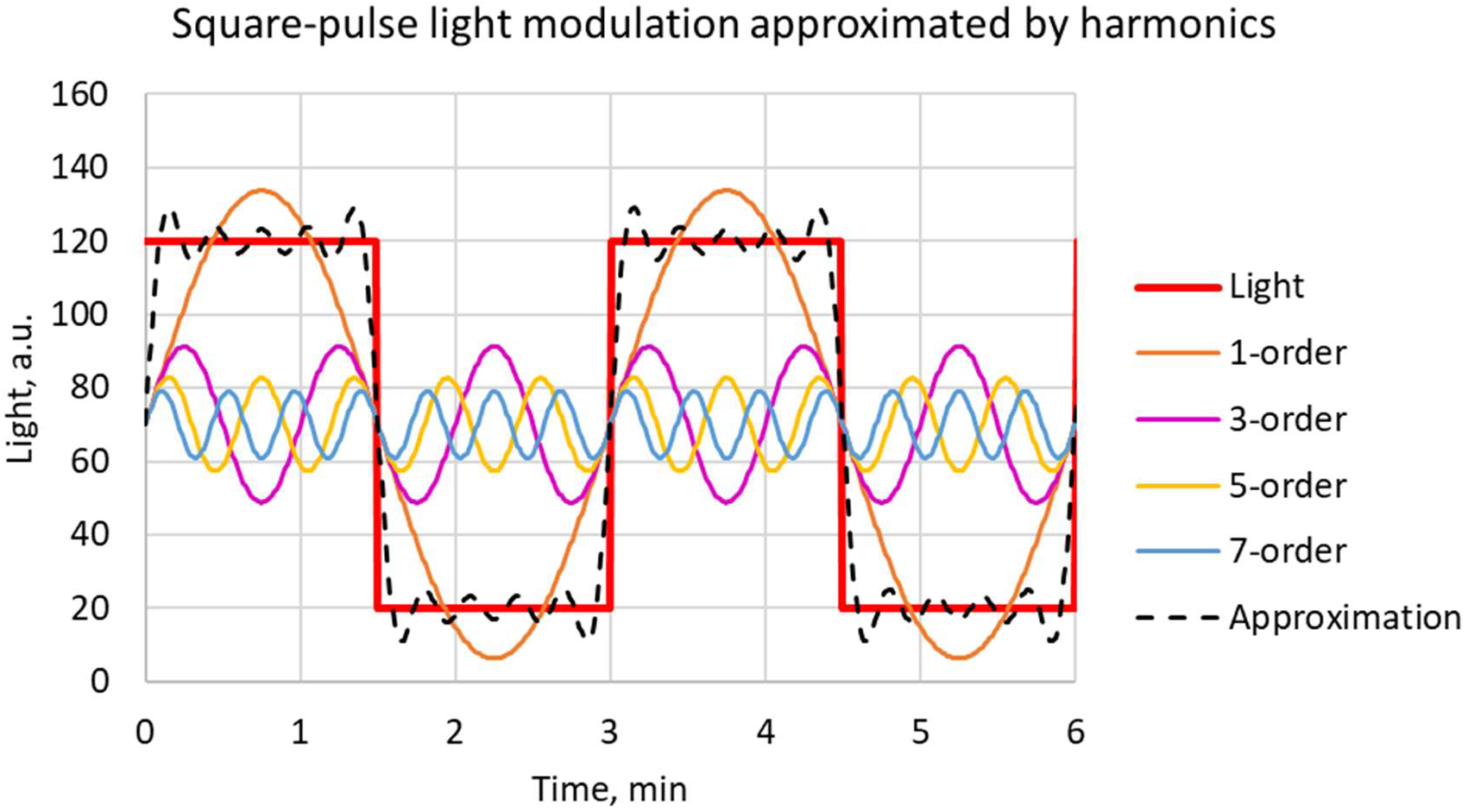
An approximation (dashed black line) of a square-modulated light (thick red line) by four harmonic components. The thin colored lines show the contributing harmonic components with periods 3 min (orange, 1^st^ -order), 1 min (magenta, 3^rd^ -order), 3/5 min (yellow, 5^th^ -order), and 3/7 min (blue, 7^th^ -order).

### SI Raw ChlF-yield data

The periods of light oscillations were changing from 8 min to 1 s (green line in Fig.SI-4). The first three oscillations had a period of 8 minutes. The ChlF pattern in the first period was strongly marked by the influence of the pre-illumination in constant light and only the two later 8 min periods were used for the data analysis. Similarly in all shorter periods (4 min to 1 s), the first two oscillations were influenced by the change in the period and not analyzed. The last three oscillations representing the given period were analyzed. Some of the analyzed time intervals are marked in Fig.SI-4 by the dashed squares.

**Fig.SI-4.**
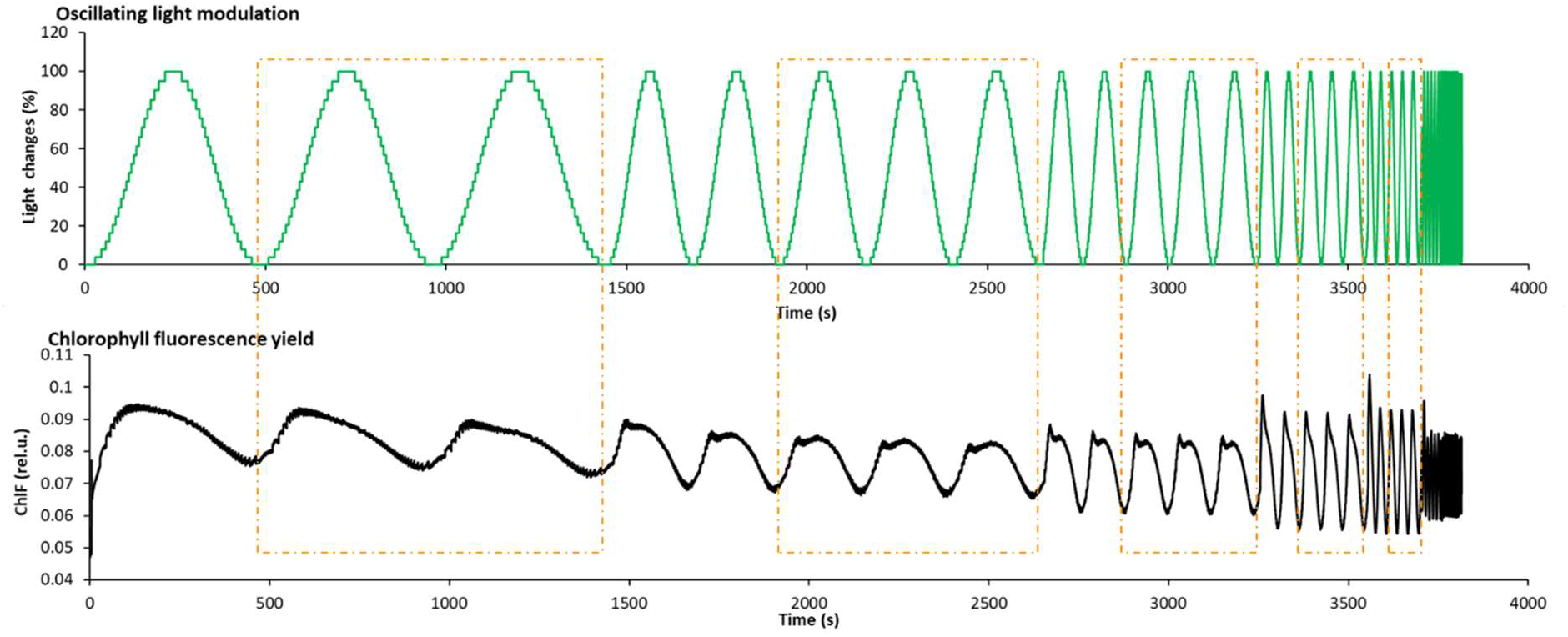
Light (green line) and ChlF yield response (black line) obtained with the protocol described in Materials and methods. The orange dashed squares mark the later periods of 8min, 4min, 2min, 60s, 30s oscillations that were used for data analysis. Cycle selection in 10s, 5s and 1s /periods is not shown, but was done in the same way.

### SI ChlF-yield data shown without separating the oscillatory and stationary components

The oscillatory part of the ChlF yield is represented in detail in Fig.3. Below, we show also the total Chl-F yield as measured in our experiments, i.e., without separating the stationary and oscillatory components.

**Fig.SI-5.**
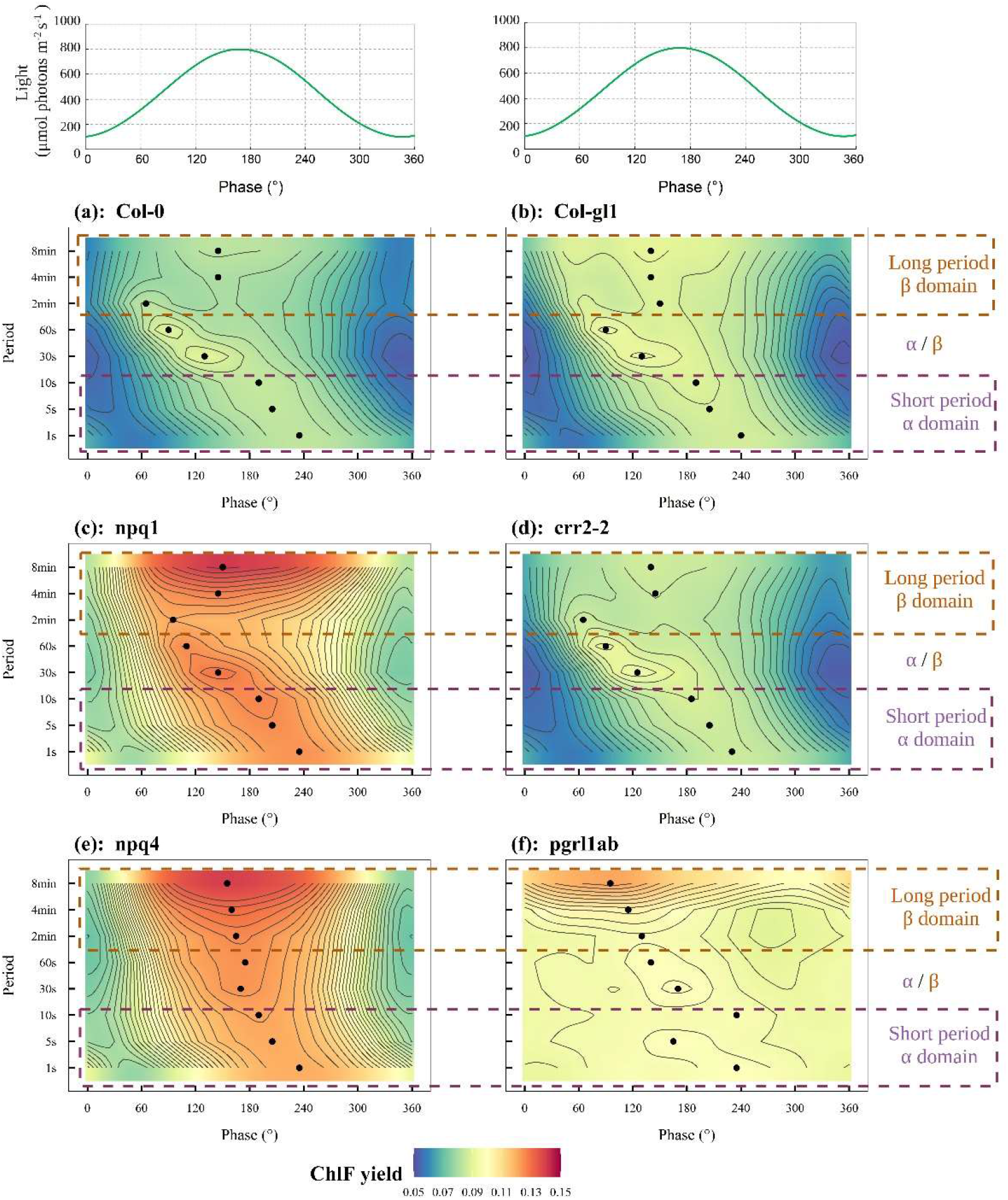
The dynamics of the total chlorophyll fluorescence (ChlF) yield (including the stationary and oscillatory components). The green line in the top panel represents light that was oscillating between 100 and 800 μmol photons·m^-2^-s^-1^ with periods in the range of 1 s to 8 min. The progress of the light oscillation and of ChlF response are shown by the phase between 0 and 360°, so that the abscissa is the same for all periods of oscillations that were applied. The other six panels show the oscillatory part of the ChlF yield induced in 6 genotypes of *A. thaliana* (n=3): (a) WT Col-0, (b) WT Col-gl1, (c) *npq1,* (d) *crr2-2,* (e) *npq4,* (f) *pgrl1ab.* The black dots indicate the phase at which maximum ChlF yield at the given period occurred. The colors ranging from blue to red represent the amplitude of ChlF signals from low to high. The contour lines separate signal ranges of 0.0025 of relative units. The brown dashed rectangles indicate long periods of 2, 4 and 8 min (β domain), and the purple dashed rectangles indicate short periods of 1, 5, and 10 s (α domain).

### SI Influence of light history on the signal patterns

The pre-illumination of the measured leaves by constant light and the changes in the period of light oscillations influenced the dynamic patterns that we measured during the first oscillations following the respective transition (Fig.SI-4). The shape of the signal patterns then converged to a largely stationary pattern in the successive two or three cycles of the same frequency, marked by the dashed squares in Fig.SI 4, that were then analyzed.

This convergence does not exclude however that the signal patterns change over longer periods of time and that the signals would be, e.g., dependent on the order in which the periods changed. This was checked in the experiments shown in Figs.SI-6 to 9 that compared signals recorded with the periods decreasing from 8 min to 1 s with those obtained for the periods increasing from 1 s to 8 min.

**Fig.SI-6.**
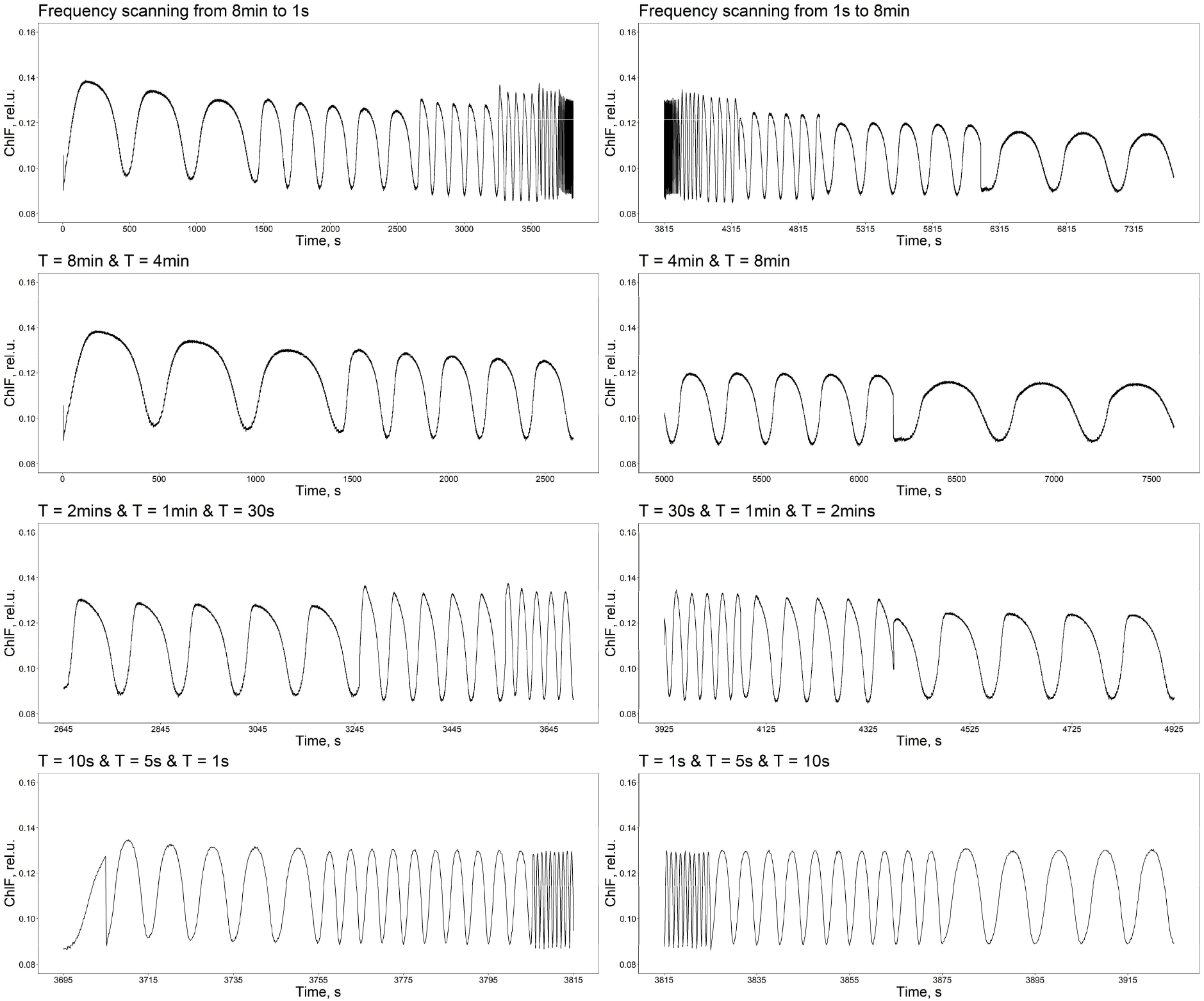
Effects of the period-scanning direction on the Chl-F responses. The top-left panel shows the ChlF signals obtained with the protocol described in Materials and Methods, i.e., with periods decreasing from 8 min to 1 s. The top-right panel depicts the ChlF signals responding to periods /changing in the opposite direction, i.e., increasing 1 s to 8 min. The second, third, and fourth rows show the comparisons in detail. The data show responses of the *npq1* mutant. Similar tests were performed with the other genotypes used in this study.

**Fig.SI-7.**
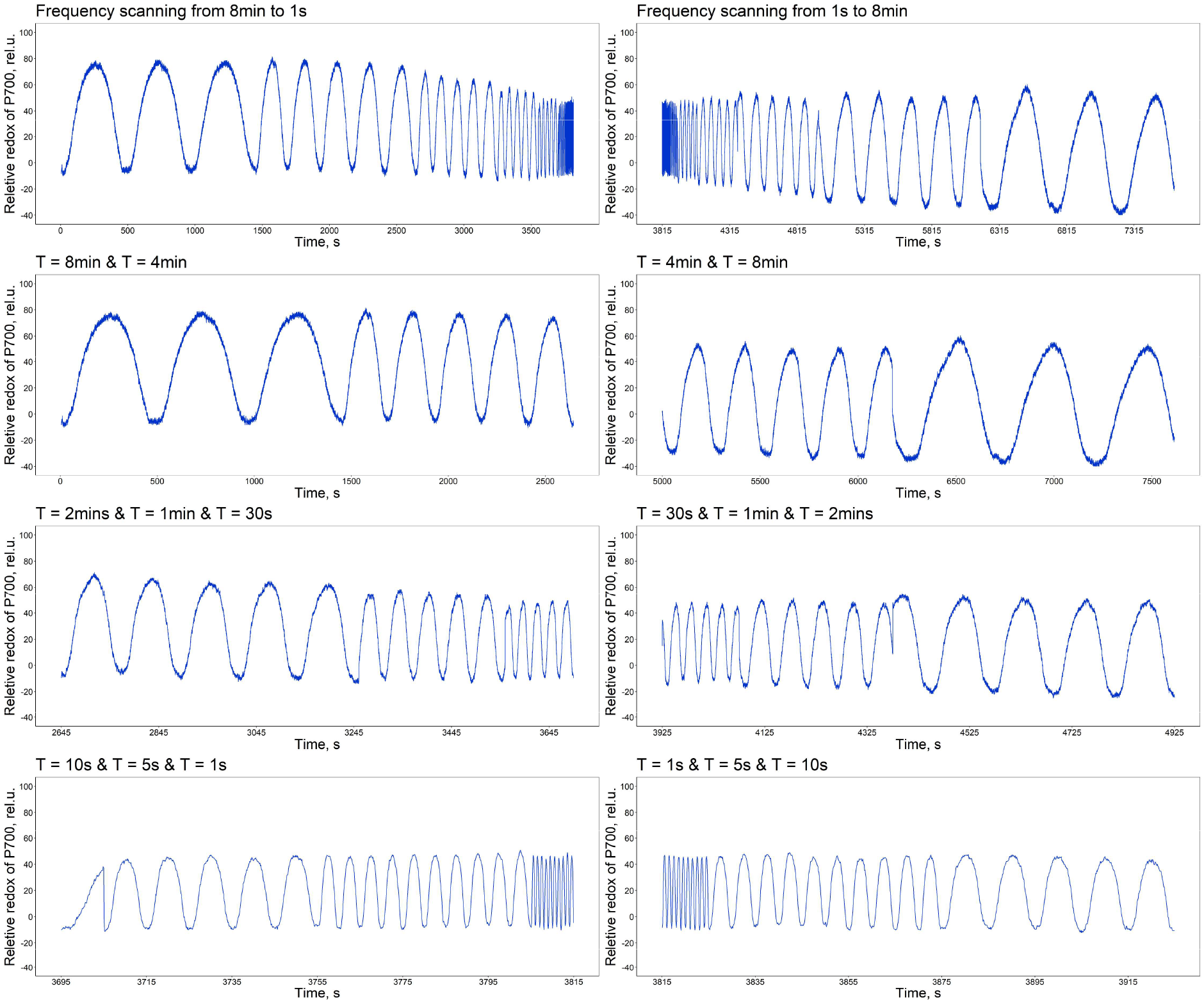
Effects of the period-scanning direction on the relative P700 redox responses. The top-left panel shows the P700 redox signals obtained with the protocol described in Materials and Methods, i.e., with periods decreasing from 8 min to 1 s. The top-right panel depicts the P700 redox signals responding to periods /changing in the opposite direction, i.e., increasing 1 s to 8 min. The second, third, and fourth rows show the comparisons in detail. The data show responses of the *npq1* mutant. Similar tests were performed with the other genotypes used in this study.

**Fig.SI-8.**
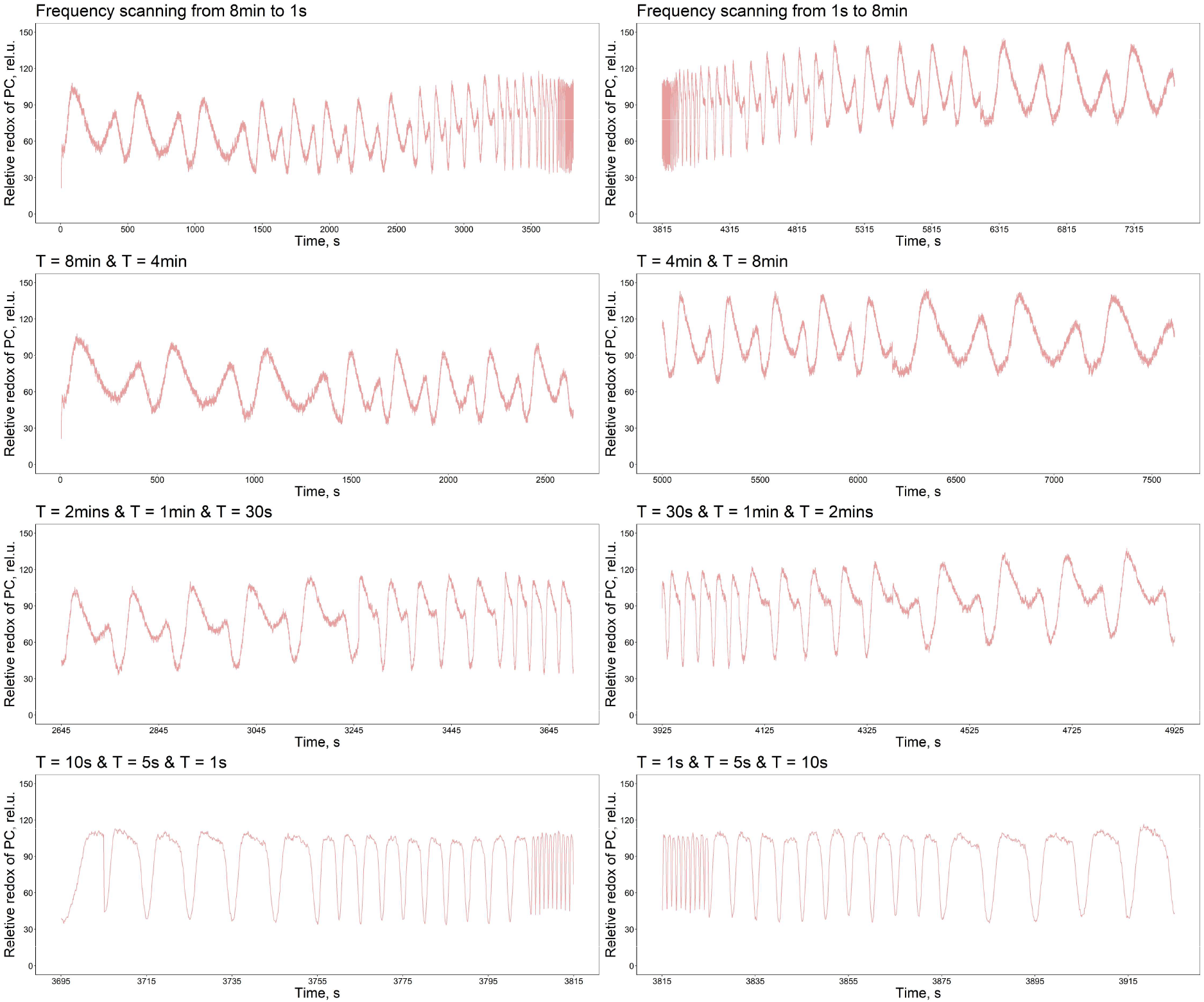
Effects of the period-scanning direction on the PC redox responses. The top-left panel shows the PC redox signals obtained with the protocol described in Materials and Methods, i.e., with periods decreasing from 8 min to 1 s. The top-right panel depicts the PC redox signals responding to periods /changing in the opposite direction, i.e., increasing 1 s to 8 min. The second, third, and fourth rows show the comparisons in detail. The data show responses of the *npq1* mutant. Similar tests were performed with the other genotypes used in this study.

**Fig.SI-9.**
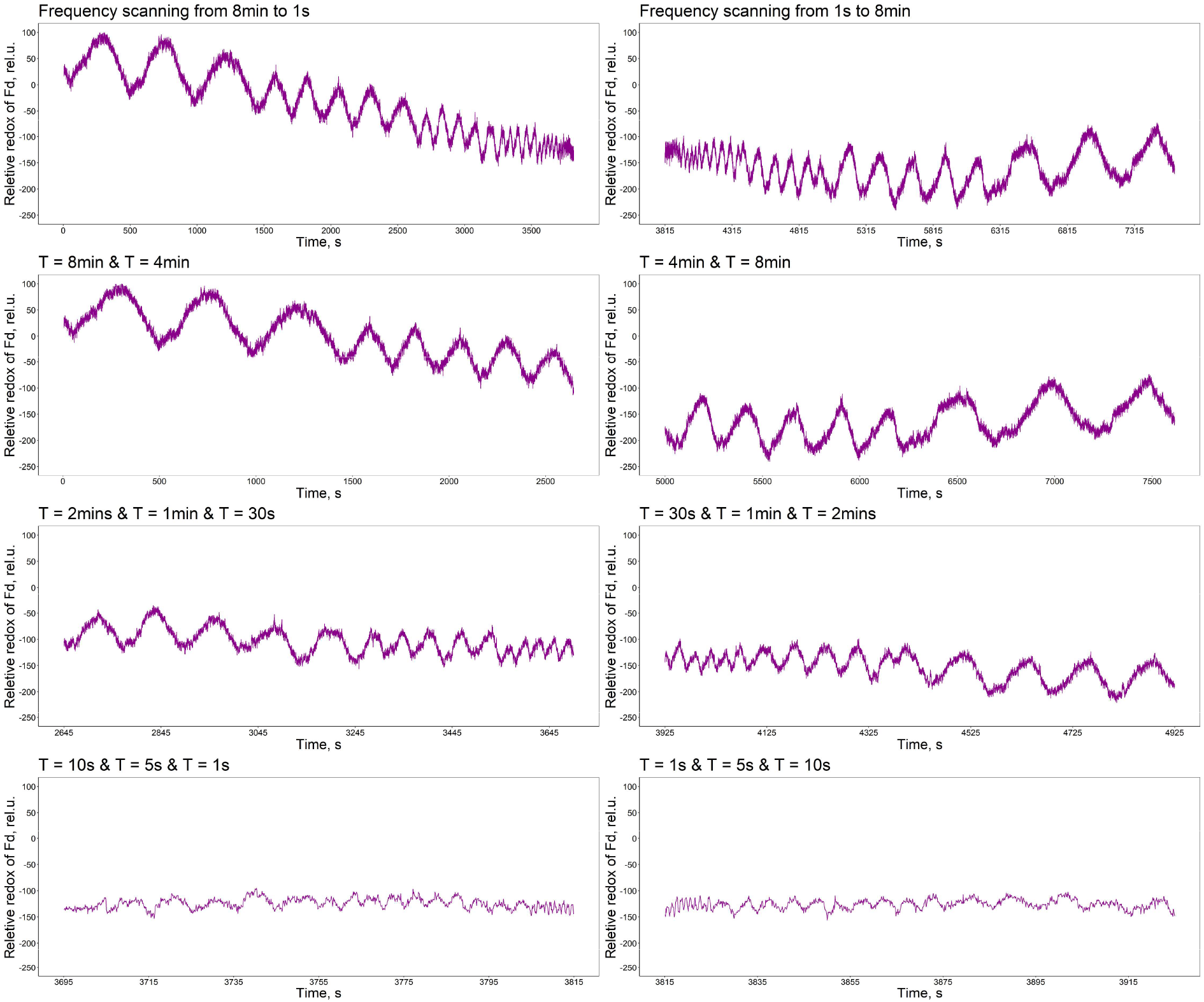
Effects of the period-scanning direction on the Fd redox responses. The top-left panel shows the Fd redox signals obtained with the protocol described in Materials and Methods, i.e., with periods decreasing from 8 min to 1 s. The top-right panel depicts the Fd redox signals responding to periods /changing in the opposite direction, i.e., increasing 1 s to 8 min. The second, third, and fourth rows show the comparisons in detail. The data show responses of the *npq1* mutant. Similar tests were performed with the other genotypes used in this study.

## References

Chazdon RL, Pearcy RW. 1991. The importance of sunflecks for forest understory plants - photosynthetic machinery appears adapted to brief, unpredictable periods of radiation. BioScience 41(11): 760–766.

Colombo M, Suorsa M, Rossi F, Ferrari R, Tadini L, Barbato R, Pesaresi P. 2016. Photosynthesis control: An underrated short-term regulatory mechanism essential for plant viability. Plant signaling & behavior 11(4): e1165382.

DalCorso G, Pesaresi P, Masiero S, Aseeva E, Nemann DS, Finazzi G, Joliot P, Barbato R, Leister D. 2008. A complex containing PGRL1 and PGR5 is involved in the switch between linear and cyclic electron flow in Arabidopsis. Cell 132(2): 273–285.

Delieu TJ, Walker DA. 1983. Simultaneous measurement of oxygen evolution and chlorophyll fluorescence from leaf pieces. Plant Physiology 73(3): 534–541.

Demmig-Adams B, Adams WW, Garab G, Govindjee. 2014. Non-photochemical quenching and energy dissipation in plants, algae and cyanobacteria preface: Springer Netherlands.

Demmig B, Winter K, Kruger A, Czygan FC. 1987. Photoinhibition and zeaxanthin formation in intact leaves - a possible role of the xanthophyll cycle in the dissipation of excess light energy. Plant Physiology 84(2): 218–224.

Ganusov VV. 2016. Strong inference in mathematical modeling: a method for robust science in the twenty-first Century. Frontiers in Microbiology 7: 1131.

Gilmore AM. 1997. Mechanistic aspects of xanthophyll cycle-dependent photoprotection in higher plant chloroplasts and leaves. Physiologia Plantarum 99(1): 197–209.

Graham PJ, Nguyen B, Burdyny T, Sinton D. 2017. A penalty on photosynthetic growth in fluctuating light. Scientific Reports 7(1): 12513.

Hashimoto M, Endo T, Peltier G, Tasaka M, Shikanai T. 2003. A nucleus-encoded factor, CRR2, is essential for the expression of chloroplast ndhB in Arabidopsis. Plant Journal 36(4): 541–549.

Hepworth C, Wood WHJ, Emrich-Mills TZ, Proctor MS, Casson S, Johnson MP. 2021. Dynamic thylakoid stacking and state transitions work synergistically to avoid acceptor-side limitation of photosystem I. Nature Plants 7(1): 87–98.

Hertle AP, Blunder T, Wunder T, Pesaresi P, Pribil M, Armbruster U, Leister D. 2013. PGRL1 is the elusive ferredoxin-plastoquinone reductase in photosynthetic cyclic electron flow. Molecular cell 49(3): 511–523.

Höhner R, Pribil M, Herbstová M, Lopez LS, Kunz HH, Li M, Wood M, Svoboda V, Puthiyaveetil S, Leister D, et al. 2020. Plastocyanin is the long-range electron carrier between photosystem II and photosystem I in plants. Proceedings of the National Academy of Sciences of the United States of America 117(26): 15354–15362.

Holzwarth AR, Miloslavina Y, Nilkens M, Jahns P. 2009. Identification of two quenching sites active in the regulation of photosynthetic light-harvesting studied by time-resolved fluorescence. Chemical Physics Letters 483(4-6): 262–267.

Horton P, Ruban A. 2005. Molecular design of the photosystem II light-harvesting antenna: photosynthesis and photoprotection. Journal of Experimental Botany 56(411): 365–373.

Jahns P, Holzwarth AR. 2012. The role of the xanthophyll cycle and of lutein in photoprotection of photosystem II. Biochimica et Biophysica Acta - Bioenergetics 1817(1): 182–193.

Johnson JE, Berry JA. 2021. The role of Cytochrome b6f in the control of steady-state photosynthesis: a conceptual and quantitative model. Photosynthesis Research 148(3): 101–136.

Johnson MP, Wientjes E. 2020. The relevance of dynamic thylakoid organisation to photosynthetic regulation. Biochimica et Biophysica Acta - Bioenergetics 1861(4): 148039.

Joliot P, Lavergne J, Béal D. 1992. Plastoquinone compartmentation in chloroplasts. I. Evidence for domains with different rates of photo-reduction. Biochimica et Biophysica Acta - Bioenergetics 1101(1): 1–12.

Kaiser E, Morales A, Harbinson J. 2018. Fluctuating light takes crop photosynthesis on a rollercoaster ride. Plant Physiology 176(2): 977–989.

Kaiser E, Morales A, Harbinson J, Heuvelink E, Prinzenberg AE, Marcelis LF. 2016. Metabolic and diffusional limitations of photosynthesis in fluctuating irradiance in Arabidopsis thaliana. Scientific Reports 6: 31252.

Kihara M, Ushijima T, Yamagata Y, Tsuruda Y, Higa T, Abiko T, Kubo T, Wada M, Suetsugu N, Gotoh E. 2020. Light-induced chloroplast movements in Oryza species. Journal of Plant Research 133(4): 525–535.

Kirchhoff H, Hall C, Wood M, Herbstová M, Tsabari O, Nevo R, Charuvi D, Shimoni E, Reich Z. 2011. Dynamic control of protein diffusion within the granal thylakoid lumen. Proceedings of the National Academy of Sciences of the United States of America 108(50): 20248–20253.

Kirchhoff H, Horstmann S, Weis E. 2000. Control of the photosynthetic electron transport by PQ diffusion microdomains in thylakoids of higher plants. Biochimica et Biophysica Acta - Bioenergetics 1459(1): 148–168.

Kitano H. 2001. Foundations of systems biology: The MIT Press Cambridge, Massachusetts London, England.

Klughammer C, Schreiber U. 2016. Deconvolution of ferredoxin, plastocyanin, and P700 transmittance changes in intact leaves with a new type of kinetic LED array spectrophotometer. Photosynthesis Research 128(2): 195–214.

Knapp AK, Smith WK. 1987. Stomatal and photosynthetic responses during sun/shade transitions in subalpine plants: influence on water use efficiency. Oecologia 74(1): 62–67.

Kono M, Noguchi K, Terashima I. 2014. Roles of the cyclic electron flow around PSI (CEF-PSI) and O2-dependent alternative pathways in regulation of the photosynthetic electron flow in short-term fluctuating light in Arabidopsis thaliana. Plant and Cell Physiology 55(5): 990–1004.

Kono M, Terashima I. 2014. Long-term and short-term responses of the photosynthetic electron transport to fluctuating light. Journal of Photochemistry and Photobiology B: Biology 137: 89–99.

Kouřil R, Strouhal O, Nosek L, Lenobel R, Chamrád I, Boekema EJ, Šebela M, Ilík P. 2014. Structural characterization of a plant photosystem I and NAD(P)H dehydrogenase supercomplex. Plant Journal 77(4): 568–576.

Külheim C, Ågren J, Jansson S. 2002. Rapid regulation of light harvesting and plant fitness in the field. Science 297(5578): 91–93.

Laughlin TG, Savage DF, Davies KM. 2020. Recent advances on the structure and function of NDH-1: The complex I of oxygenic photosynthesis. Biochimica et Biophysica Acta - Bioenergetics 1861(11): 148254–148254.

Lazár D, Kaňa R, Klinkovský T, Nauš J. 2005. Experimental and theoretical study on high temperature induced changes in chlorophyll a fluorescence oscillations in barley leaves upon 2% CO2. Photosynthetica 43(1): 13–27.

Lazár D, Niu Y, Nedbal L. 2022. Insights on the regulation of photosynthesis in pea leaves exposed to oscillating light. Journal of Experimental Botany 73(18): 6380–6393.

Li M, Mukhopadhyay R, Svoboda V, Oung HMO, Mullendore DL, Kirchhoff H. 2020. Measuring the dynamic response of the thylakoid architecture in plant leaves by electron microscopy. Plant Direct 4(11): e00280.

Li TY, Shi Q, Sun H, Yue M, Zhang SB, Huang W. 2021. Diurnal response of photosystem I to fluctuating light is affected by stomatal conductance. Cells 10(11): 3128.

Li X-P, Björkman O, Shih C, Grossman AR, Rosenquist M, Jansson S, Niyogi KK. 2000. A pigment-binding protein essential for regulation of photosynthetic light harvesting. Nature 403(6768): 391–395.

Li X-P, Gilmore AM, Niyogi KK. 2002. Molecular and global time-resolved analysis of a psbS gene dosage effect on pH-and xanthophyll cycle-dependent nonphotochemical quenching in photosystem II. Journal of Biological Chemistry 277(37): 33590–33597.

Long SP, Taylor SH, Burgess SJ, Carmo-Silva E, Lawson T, De Souza AP, Leonelli L, Wang Y. 2022. Into the shadows and back into sunlight: photosynthesis in fluctuating light. Annual Review of Plant Biology 73(1): 617–648.

Malone LA, Proctor MS, Hitchcock A, Hunter CN, Johnson MP. 2021. Cytochrome b6f – Orchestrator of photosynthetic electron transfer. Biochimica et Biophysica Acta - Bioenergetics 1862(5): 148380.

Matthews JSA, Vialet-Chabrand S, Lawson T. 2018. Acclimation to fluctuating light impacts the rapidity of response and diurnal rhythm of stomatal conductance. Plant Physiology 176(3): 1939–1951.

Mitchell-Olds T. 2001. Arabidopsis thaliana and its wild relatives: a model system for ecology and evolution. Trends in Ecology & Evolution 16(12): 693–700.

Muller P, Li XP, Niyogi KK. 2001. Non-photochemical quenching. A response to excess light energy. Plant Physiology 125(4): 1558–1566.

Munekage Y, Hojo M, Meurer J, Endo T, Tasaka M, Shikanai T. 2002. PGR5 is involved in cyclic electron flow around photosystem I and is essential for photoprotection in Arabidopsis. Cell 110(3): 361–371.

Murchie EH, Ruban AV. 2020. Dynamic non-photochemical quenching in plants: from molecular mechanism to productivity. Plant Journal 101(4): 885–896.

Nakano H, Yamamoto H, Shikanai T. 2019. Contribution of NDH-dependent cyclic electron transport around photosystem I to the generation of proton motive force in the weak mutant allele of pgr5. Biochimica et Biophysica Acta - Bioenergetics 1860(5): 369–374.

Nedbal L, Březina V. 2002. Complex metabolic oscillations in plants forced by harmonic irradiance. Biophysical Journal 83(4): 2180–2189.

Nedbal L, Březina V, Červený J, Trtílek M. 2005. Photosynthesis in dynamic light: systems biology of unconventional chlorophyll fluorescence transients in Synechocystis sp. PCC 6803. Photosynthesis Research 84(1): 99–106.

Nedbal L, Lazár D. 2021. Photosynthesis dynamics and regulation sensed in the frequency domain. Plant Physiology 187(2): 646–661.

Nilkens M, Kress E, Lambrev P, Miloslavina Y, Muller M, Holzwarth AR, Jahns P. 2010. Identification of a slowly inducible zeaxanthin-dependent component of non-photochemical quenching of chlorophyll fluorescence generated under steady-state conditions in Arabidopsis. Biochimica et Biophysica Acta - Bioenergetics 1797(4): 466–475.

Niyogi KK, Grossman AR, Björkman O. 1998. Arabidopsis mutants define a central role for the xanthophyll cycle in the regulation of photosynthetic energy conversion. The Plant Cell 10(7): 1121–1134.

Ogata K. 2010.Modern control engineering. Boston: Prentice-Hall.

Pantazopoulou CK, Bongers FJ, Pierik R. 2021. Reducing shade avoidance can improve Arabidopsis canopy performance against competitors. Plant, Cell & Environment 44(4): 1130–1141.

Peltier G, Aro EM, Shikanai T. 2016. NDH-1 and NDH-2 plastoquinone reductases in oxygenic photosynthesis. Annual Review of Plant Biology 67: 55–80.

Peressotti A, Marchiol L, Zerbi G. 2001. Photosynthetic photon flux density and sunfleck regime within canopies of wheat, sunflower and maize in different wind conditions. Italian Journal of Agronomy 4: 87–92.

Pintelon R, Schoukens J. 2012. System identification: a frequency domain approach: John Wiley & Sons.

Pospíšil P. 2009. Production of reactive oxygen species by photosystem II. Biochimica et Biophysica Acta - Bioenergetics 1787(10): 1151–1160.

Roach T, Krieger-Liszkay A. 2012. The role of the PsbS protein in the protection of photosystems I and II against high light in Arabidopsis thaliana. Biochimica et Biophysica Acta - Bioenergetics 1817(12): 2158–2165.

Ruban AV. 2016. Nonphotochemical chlorophyll fluorescence quenching: mechanism and effectiveness in protecting plants from photodamage. Plant Physiology 170(4): 1903–1916.

Ruban AV, Johnson MP. 2015. Visualizing the dynamic structure of the plant photosynthetic membrane. Nature Plants 1(11): 15161.

Schreiber U, Klughammer C. 2016. Analysis of photosystem I donor and acceptor sides with a new type of online-deconvoluting kinetic LED-array spectrophotometer. Plant and Cell Physiology 57(7): 1454–1467.

Schwartz L. 2008. Mathematics for the physical sciences: Courier Dover Publications.

Sétif P, Boussac A, Krieger-Liszkay A. 2019. Near-infrared in vitro measurements of photosystem I cofactors and electron-transfer partners with a recently developed spectrophotometer. Photosynthesis Research 142(3): 307–319.

Shikanai T. 2014. Central role of cyclic electron transport around photosystem I in the regulation of photosynthesis. Current Opinion in Biotechnology 26: 25–30.

Shimakawa G, Miyake C. 2018. Changing frequency of fluctuating light reveals the molecular mechanism for P700 oxidation in plant leaves. Plant Direct 2(7): e00073.

Smith WK, Berry ZC. 2013. Sunflecks? Tree Physiology 33(3): 233–237.

Strand DD, Fisher N, Kramer DM. 2017. The higher plant plastid NAD(P)H dehydrogenase-like complex (NDH) is a high efficiency proton pump that increases ATP production by cyclic electron flow. The Journal of biological chemistry 292(28): 11850–11860.

Sugimoto K, Okegawa Y, Tohri A, Long TA, Covert SF, Hisabori T, Shikanai T. 2013. A single amino acid alteration in PGR5 confers resistance to antimycin A in cyclic electron transport around PSI. Plant and Cell Physiology 54(9): 1525–1534.

Suorsa M, Grieco M, Jarvi S, Gollan PJ, Kangasjarvi S, Tikkanen M, Aro EM. 2013. PGR5 ensures photosynthetic control to safeguard photosystem I under fluctuating light conditions. Plant signaling & behavior 8(1): e22741.

Suorsa M, Järvi S, Grieco M, Nurmi M, Pietrzykowska M, Rantala M, Kangasjärvi S, Paakkarinen V, Tikkanen M, Jansson S, et al. 2012. PROTON GRADIENT REGULATION5 is essential for proper acclimation of Arabidopsis photosystem I to naturally and artificially fluctuating light conditions. Plant Cell 24(7): 2934–2948.

Suorsa M, Rossi F, Tadini L, Labs M, Colombo M, Jahns P, Kater Martin M, Leister D, Finazzi G, Aro E-M, et al. 2016. PGR5-PGRL1-dependent cyclic electron transport modulates linear electron transport rate in Arabidopsis thaliana. Molecular Plant 9(2): 271–288.

Thormählen I, Zupok A, Rescher J, Leger J, Weissenberger S, Groysman J, Orwat A, Chatel-Innocenti G, Issakidis-Bourguet E, Armbruster U, et al. 2017. Thioredoxins play a crucial role in dynamic acclimation of photosynthesis in fluctuating light. Molecular Plant 10(1): 168–182.

Ünnep R, Zsiros O, Hörcsik Z, Markó M, Jajoo A, Kohlbrecher J, Garab G, Nagy G. 2017. Low-pH induced reversible reorganizations of chloroplast thylakoid membranes — as revealed by small-angle neutron scattering. Biochimica et Biophysica Acta - Bioenergetics 1858(5): 360–365.

von Bismarck T, Korkmaz K, Ruß J, Skurk K, Kaiser E, Galvis VC, Cruz J, Strand D, Köhl K, Eirich J, et al. 2021. Light acclimation interacts with thylakoid ion transport to govern the dynamics of photosynthesis: New Phytologist, in press.

Wada S, Amako K, Miyake C. 2021. Identification of a novel mutation exacerbated the PSI photoinhibition in pgr5/pgrl1 mutants; caution for overestimation of the phenotypes in Arabidopsis pgr5-1 mutant. Cells 10(11).

Wang CJ, Yamamoto H, Shikanai T. 2015. Role of cyclic electron transport around photosystem I in regulating proton motive force. Biochimica et Biophysica Acta - Bioenergetics 1847(9): 931–938.

Ware MA, Belgio E, Ruban AV. 2015. Comparison of the protective effectiveness of NPQ in Arabidopsis plants deficient in PsbS protein and zeaxanthin. Journal of Experimental Botany 66(5): 1259–1270.

Way DA, Pearcy RW. 2012. Sunflecks in trees and forests: from photosynthetic physiology to global change biology. Tree Physiology 32(9): 1066–1081.

Wehner A, Storf S, Jahns P, Schmid VHR. 2004. De-epoxidation of violaxanthin in light-harvesting complex I proteins. Journal of Biological Chemistry 279(26): 26823–26829.

Wood WHJ, MacGregor-Chatwin C, Barnett SFH, Mayneord GE, Huang X, Hobbs JK, Hunter CN, Johnson MP. 2018. Dynamic thylakoid stacking regulates the balance between linear and cyclic photosynthetic electron transfer. Nature Plants 4(2): 116–127.

Yamamoto H, Peng L, Fukao Y, Shikanai T. 2011. An Src homology 3 domain-like fold protein forms a ferredoxin binding site for the chloroplast NADH dehydrogenase-like complex in Arabidopsis. Plant Cell 23(4): 1480–1493.

Yamamoto H, Shikanai T. 2019. PGR5-dependent cyclic electron flow protects photosystem I under fluctuating light at donor and acceptor Sides. Plant Physiology 179(2): 588–600.

Yamori W, Makino A, Shikanai T. 2016. A physiological role of cyclic electron transport around photosystem I in sustaining photosynthesis under fluctuating light in rice. Scientific Reports 6: 20147.

## SI - References

Ferimazova N, Küpper H, Nedbal L, Trtílek M. 2002. New insights into photosynthetic oscillations revealed by two-dimensional microscopic measurements of chlorophyll fluorescence kinetics in intact leaves and isolated protoplasts. Photochemistry and Photobiology 76(5): 501–508.

Schwartz L. 2008. Mathematics for the physical sciences: Courier Dover Publications.

